# *Agouti* integrates environmental information to regulate natural variation in paternal behavior

**DOI:** 10.1101/2025.06.17.658598

**Authors:** Forrest Dylan Rogers, Sara A. Mereby, Anna M. Kasper, Sehee Kim, Ricardo Mallarino, Catherine Jensen Peña

## Abstract

Male investment in offspring rearing through paternal care is rare among mammals and the neural mechanisms governing its emergence are poorly understood. We leveraged the natural paternal behavior of African striped mice (*Rhabdomys pumilio*) in combination with brain-wide cFos quantification, single-nucleus RNA-sequencing, viral-mediated gene manipulation, and environmental manipulation to dissect the neural basis of natural variation in male parenting. We find that socio-environmental conditions drive individual variation in male alloparenting such that post-weaning social isolation increases paternal care while social living in higher density groups increases infanticide. This natural variation in care corresponds to neural activity in the medial preoptic area and changes in correlated activity across brain regions. Within the medial preoptic area, expression of agouti signaling protein (*Agouti*) in neurons is increased by group housing and is negatively associated with care, and overexpression of Agouti reduces care and enhances infanticide in previously tolerant animals. Naturalistic manipulations further reveal that *Agouti* integrates long-term housing conditions rather than food availability/hunger. Together, our results demonstrate that *Agouti* acts as a molecular integrator of socio-environmental information to drive variation in paternal care.

## INTRODUCTION

Active parental care is crucial for the survival and development of altricial offspring, with maternal care forming the evolutionary foundation of caregiving across all mammals. In all species, mothers provide essential resources such as nutrition, thermoregulation, and protection^1^. In a substantially smaller fraction of mammals (3–5%), including humans^2,3^, fathers also contribute to offspring care. While the neural circuits underpinning maternal care have been extensively studied and are conserved across mammals, our understanding of the causal neurobiological and molecular mechanisms that give rise to naturally occurring, active care in males is still in its infancy^3^.

Examination of neuroendocrine and peptidergic pathways of known importance in lactating mothers, infanticidal behavior in standard laboratory mice (*Mus musculus*) and rats (*Rattus norvegicus*), and genetic variation among deer mice and vole species have yielded insights into the paternal brain^3–5^. It is well-established that maternal care is mediated by the medial preoptic area of the hypothalamus (MPOA), a brain region that undergoes dynamic hormonal and epigenetic remodeling during pregnancy^3^. This remodeling reshapes neural circuits, suppressing pup-directed aggression whilst promoting approach and engaging downstream reward systems to reinforce caregiving^6,7^. In contrast, male laboratory mice and rats do not naturally exhibit parental care^8,9^. Following mating, they temporarily reduce infanticidal behavior10; yet merely tolerating pups does not confer the same developmental and evolutionary advantages as active caregiving.

Experimental studies have shown that male laboratory mice and rats can be induced to transiently display parental behaviors through sensitization^11^, a process typically achieved via cohabitation with a female after mating or by prolonged pup exposure. In sensitized males, the MPOA has been identified as a critical hub^12–14^, integrating upstream olfactory inputs^13,14^ and engaging downstream reward pathways^15,16^, suggesting that the fundamental neural circuitry for caregiving exists in males but remains inactive under normal conditions. However, certain mechanisms that promote maternal care appear to be absent or functionally distinct in males^17^, indicating potential sex specific cellular and/or molecular regulatory pathways. While these studies provide insight into how paternal behaviors can be elicited, they also underscore the limitations of traditional rodent models in understanding the neurobiological basis of naturally occurring variation in male caregiving, particularly within a single species.

The African striped mouse (*Rhabdomys pumilio*) is an attractive model for investigating the neurobiology of paternal care. Unlike traditional laboratory murine rodents, African striped mice (hereafter, ‘striped mice’) naturally exhibit both paternal and alloparental behaviors, with nongenetic caregivers (alloparents) readily providing care to younger siblings and even unrelated infants^18^. Striped mice thrive in a diverse range of social environments, from communal group living to philopatry, territorial breeding, and solitary wandering^18–21^. When male striped mice disperse from their natal nest, they assume one of a variety of social strategies according to prevailing environmental pressures including population density and resource availability. These strategies include solitary roaming and the formation of bachelor groups (with kin and nonkin)^20,22–24^. This ecological flexibility extends to laboratory settings, where they can be housed under conditions that mimic these social strategies. Notably, unlike laboratory rats and mice, male striped mice do not require prior reproductive experience to display parental care^19^, and their parental phenotypes exhibit natural intraspecies variation. This marked variation within a single species, where genetic factors are controlled, offers a unique opportunity to dissect the molecular and neurobiological mechanisms underlying experience-dependent individual differences in paternal care. Here, leveraging the striped mouse as a model and employing an integrative approach— including brain-wide quantification of cFos expression, single-nucleus RNA sequencing, viral-mediated gene manipulation in the brain, and socio-environmental manipulations—we provide new insight into the neural mechanisms driving naturally occurring paternal care.

## RESULTS

### Striped mouse males are primed to provide care

While previous field and laboratory studies have shown that sexually naïve male striped mice (Figure 1a) exhibit paternal behavior^18,25,26^, the full extent and quality of this care as well as how it compares to that of genetic fathers (i.e., sires) and mothers (i.e., dams) in the laboratory setting has not been fully characterized. To this end, we compared the parental behavior of laboratory-reared sexually naïve males with that of sires and dams in a pup interaction test^27^. The temporal dynamics of behavior were highly stereotyped across sex and reproductive experience (Supplemental Video 1), with nearly all animals initially crossing the enclosure to inspect (i.e., sniff) pup stimuli with a subsequent transition into either caregiving or infanticidal behaviors. Care behaviors included licking and grooming, huddling contact, and lateral contact (Supplemental Table 1; Supplemental Data Figure 1).

**Fig. 1.**
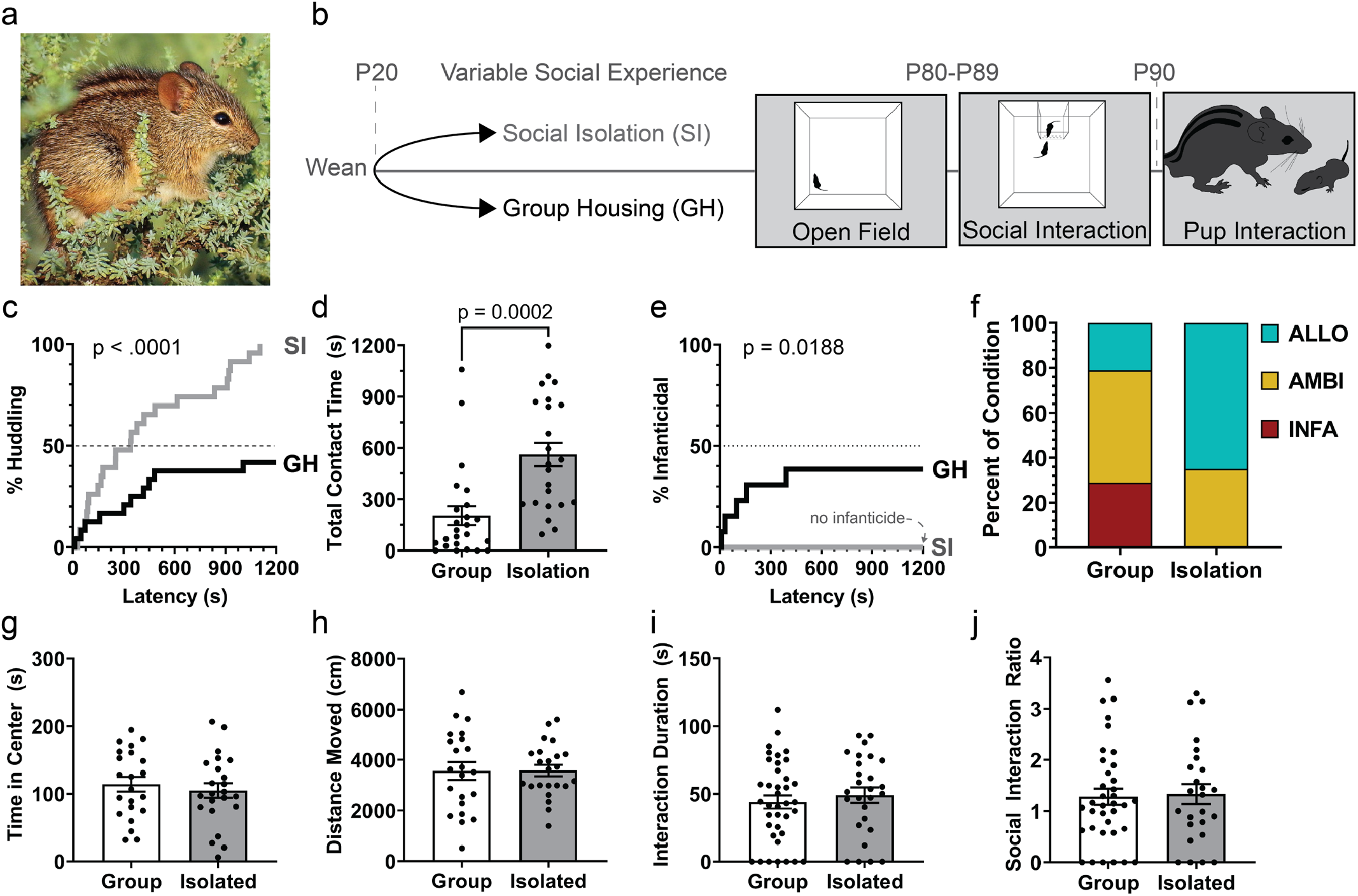
Allopaternal care in African striped mice. (a) Photograph of an African striped mouse (Photo credit: Marie Delport, iNaturalist). (b) Schematic of experimental design and behavioral testing timeline (c) Survival curve illustrating the proportion of males assuming a huddling posture over a novel pup stimulus, comparing SI (grey) and GH (black) rearing conditions (Log-rank Mantel-Cox test, X2= 16.86, df = 1, p < 0.0001). (d) Cumulative contact time (sum of huddling, licking/grooming, and adjacent contact time) with pups across rearing conditions (Unpaired t-test, t(45) = 4.063, p = 0.0002, Cohen’s d = 1.211, CI95% = [180.2, 534.3]). (e) Proportion of infanticide in males from different rearing conditions (X2= 5.521, df = 1, p = 0.0188). (f) Proportional representation of (INFA), ambivalent (AMBI), or allopaternal (ALLO) responses towards pups in males from different rearing conditions (Chi-square test on proportions: X2 = 54.16, df = 2, p < .0001). (g,h) Open field test results showing time spent in the center of the open field (g; mixed effects model: t(45.00) = -1.486, p = 0.144) and total distance traveled (h; t(43.46) = 0.574, p = 0.569). (i,j) Social interaction test results, including time interacting with an adult, same-sex conspecific (i; t(63.00) = 0.674, p = 0.503) and social interaction ratio across different conditions (j, t(63.00) = 0.392, p = 0.697). Social interaction ratio was calculated as the time spent exploring a novel mouse over time exploring an empty enclosure. Error bars represent standard error of means.

Collectively, males exhibited a wide range of phenotypes, from “dyspaternal” (pup ambivalence or infanticide) to “allopaternal.” We defined ambivalence as neglect following an initial inspection of the pup stimulus (e.g., Supplemental Video 2), while allopaternal males displayed licking, grooming, and huddling behaviors without prior reproductive experience at levels comparable to sires and dams (Supplemental Data Figure 1, Supplemental Data Table 2, Supplemental Video 3). Thus, our results confirm that reproductive experience is not a prerequisite for the display of paternal care in male striped mice.

### Social environment drives variation in male parental care

In the wild, striped mice exhibit social flexibility, naturally forming either complex social groups or living in isolation, depending on environmental factors such as season and population density^19,20,28^, and prior studies have shown that social environment (e.g., social isolation vs. familial housing) can alter male care^19^. We leveraged this social flexibility to explicitly determine how social environment influences development of male parental care behaviors by rearing males either in social isolation (SI) or group housing with 3-4 age-matched males (GH) from weaning until sexual maturity (≥ PND90) (Figure 1b). This developmental period encompasses the window in which many striped mouse males would disperse from their natal nest to either roam in isolation, find their own territory, or join a bachelor group^20,22–24,29^. In the pup interaction test, SI males engaged in significantly higher levels of all individual paternal behaviors (e.g., huddling, licking/grooming, and other pup contact) than GH males (Figure 1c-e). Social environment also shaped overall paternal phenotypes: 21% of GH males were allopaternal, while 29% were infanticidal, with the remaining ambivalent (Figure 1f). In contrast, 65% of SI males were allopaternal, and none displayed infanticide (Figure 1f).

To determine whether differences in allopaternal care could be attributed to baseline anxiety-like, exploratory, or social behaviors driven by rearing environment, we assessed males in open field and social interaction tests. SI did not alter time in the center of the open field or total distance traveled, indicating no anxiogenic or exploratory effects of SI in striped mice—contrary to findings in Mus^30^ (Figure 1g/h). Similarly, SI did not alter adult social interactions (Figure 1i/j). Together, these findings suggest that rearing environment influences paternal care without altering other exploratory or social behaviors.

### The MPOA is a hub of natural variation in male parental care

To identify brain regions implicated in paternal behavior in striped mice and determine whether neural activity in these regions is associated with individual differences, we exposed sexually naïve GH and SI striped mouse males to either pups (n=11/group) or an empty cage (n=10/group) for 60 minutes. We then quantified brain-wide cFOS expression as a marker of neural activity (Figure 2a). Regardless of rearing condition, pup-exposed animals exhibited greater cFOS than controls in the MPOA, anterior olfactory nucleus (AON), and prefrontal cortex (PFC, consisting of the infralimbic and prelimbic cortex) (Supplemental Data Figure 2, Supplemental Data Tables 3/4).

**Fig. 2.**
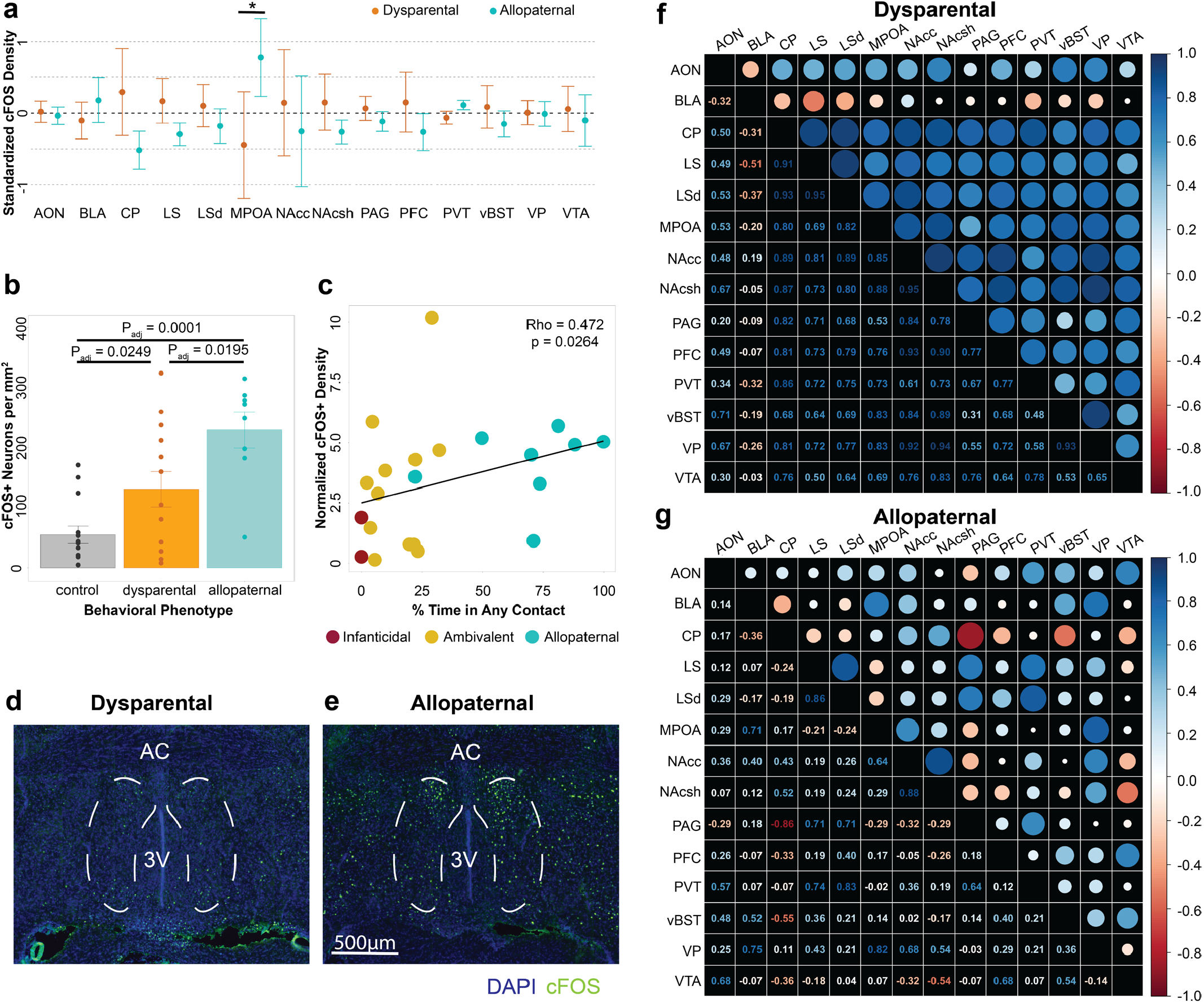
Immediate early gene expression in pup-exposed animals. (a) Brain-wide density of active (cFOS+) neurons. Dots represent group means for dysparental (orange) and allopaternal (turquoise) males. Asterix indicates Padj < .05. (b) Density of active (cFOS+) neurons within the medial preoptic area (MPOA) in sexually-naïve striped mice exposed to either a novel pup stimulus and grouped into resulting phenotypes (dysparental or allopaternal) or an empty cage (control). (c) Scatter plot depicting the correlation between the density of active MPOA neurons and the percentage of time spent in any contact (i.e., huddling, lateral, and licking/grooming). (d,e) Representative images of cFOS+ neurons in males with dysparental (d) and allopaternal (e) phenotypes. (f/g) Matrices of Spearman’s rank correlations between ROIs in (f) dysparental and (g) allopater-nal phenotypes. Circle size and color is proportional to the absolute value of the correlation. Abbreviations: accessory olfactory nucleus (AON); basolateral amygdala (BLA); caudate putamen (CP); lateral septum (LS), dorsal lateral septum (LSd); nucleus accumbens (NAc); prefrontal cortex (PFC); paraventricular nucleus of the thalamus (PVT); ventral bed nucleus of the stria terminalis (vBST); ventral pallidum (VP); ventral tegmental area (VTA). Error bars represent standard error of means.

We next examined activity by phenotype, regardless of rearing condition (n=8 allopaternal, 14 dysparental, and 20 control). Allopaternal and dysparental males had greater cFOS than controls in the AON, MPOA, PFC, and ventral tegmental area (VTA), with allopaternal males also exhibiting higher activity in the basolateral amygdala (BLA) compared to controls (Supplemental Data Figure 3, Supplemental Data Tables 5/6). The only region in which allopaternal and dyspaternal males differed was the MPOA, which showed significantly greater activity in allopaternal males (Figure 2b/d/e; Supplemental Data Table 6). Additionally, cFOS+ neuronal density (cFOS+ neurons per mm2) correlated significantly with total caring contact (rho = .47; Figure 2c) and huddling (rho = .48) exclusively in the MPOA, with no significant correlations observed in other regions of interest (Figure 2c; Supplemental Data Table 7).

Across all pup-exposedmales, MPOA activity correlated with that of nine other brain regions examined, including the PFC, paraventricular nucleus of the thalamus (PVT), dorsal lateral septum (LSd), caudate putamen (CP), ventral pallidum (VP), nucleus accumbens core (NAcc), nucleus accumbens shell (NAcsh), ventral bed nucleus of the stria terminalis (vBST), and VTA. Furthermore, the strength and significance of these correlations varied by phenotype, with greater MPOA-correlated activity in BLA and VP among paternal males (Figure 2f/g, Supplemental Data Figure 4; Supplemental Data Table 8). Together, these findings show that the MPOA is a central hub for paternal care in striped mice, supporting the hypothesis that males co-opt maternal neural circuitry, and that MPOA regional and circuit-level activity reflects individual variation in paternal care.

### Cellular composition of the striped mouse preoptic area

While cFOS activity within MPOA distinguished paternal from dyspaternal males, we next sought to determine whether cell-type specific composition and/or molecular processes within the MPOA drive these individual differences in care. To do so, we performed single-nucleus RNA-sequencing (snRNA-seq) using the 10X Genomics platform in MPOA samples from sexually naïve alloparental (n=3), infanticidal (n=3), or control males (n=3), as well as sires (n=4), and dams (n=4) (Figure 3a/b). GH males were behaviorally phenotyped, and on a subsequent day, all except controls were exposed to pup stimuli for 5 minutes before harvesting tissue to capture pup-induced gene expression response (Figure 3a). After doublet removal and quality filtering, we obtained 164,507 nuclei (Figure 3c), with each sample yielding an average of 9,677 nuclei and 1206 genes/ nucleus aligned to the *Rhabdomys pumilio* genome^31^.

**Fig. 3.**
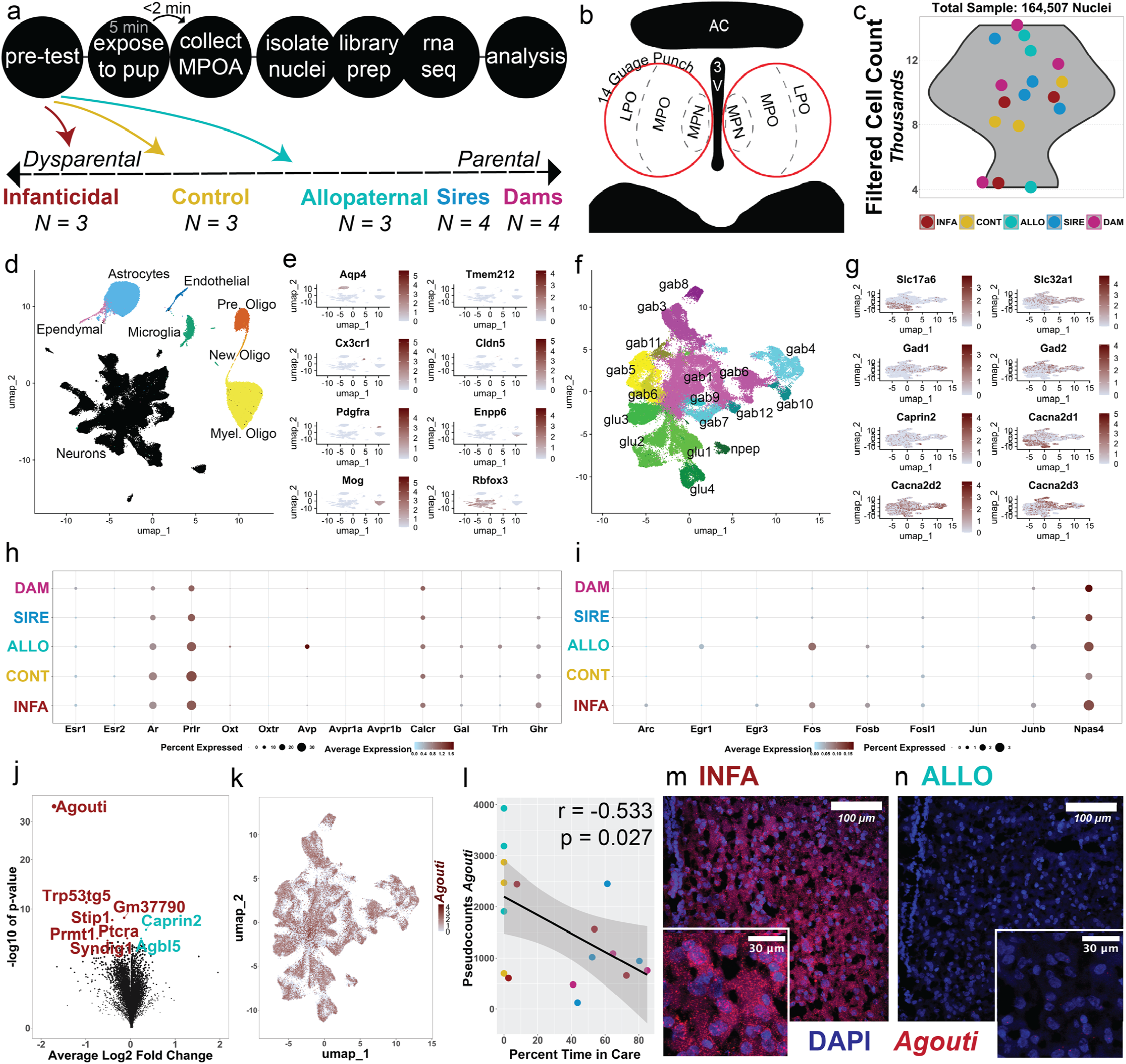
Immediate early gene expression in pup-exposed animals. (a) Schematic of experimental design and behavioral categories included in our analysis; (b) Diagram depicting the region isolated for single-nucleus RNA-sequencing. (c) Count, after quality control filtering, of sequenced nuclei for each sample. Colors correspond to behavioral category. (d,e) UMAP of clustered nuclei depicting cell classes (d) based on expression of known marker genes (e). (f,g) 17 neuronal sub-clusters segregated by release type (i.e., GABAergic, glutamatergic, and neuropeptidergic). Colors correspond to alternative sub-clusters, grouped in four subclasses of nuclei organized by voltage-dependent calcium channel subunits [i.e., Cac-na2d1 (green), Cacna2d2 (cyan), Cacna2d3 (magenta), or none (yellow)]. Expression of marker genes (g). (h) Expression of a priori neuropeptides of interest across behavioral groups. (i) Immediate early genes expression across behavioral groups. (j) DEG analysis of pseudobulked expression within all neurons with gene expression enriched in infanticidal (left, red) vs allopaternal (right, turquoise) males. Named genes were significant after FDR correction (FDR ≤ 0.1). (k, l) Agouti expression across different neuronal clusters. (l) Correlation of Agouti levels and percent time displaying care behaviors (t(15) = -2.4233, p = 0.0285, r = -0.530)). (m-n) Example RNAscope images of Agouti expression (red) in the MPOA of infanticidal (INFA; m) and allopaternal (ALLO; n) sexually naïve males.

We performed *Harmony* integration^32^, computed nearest neighbors, and identified eight major cell clusters using nearest neighbor modularity optimization, projected onto a uniform manifold and approximation projection (UMAP) embedding (Figure 3d). Clusters were classified based on known marker genes for neurons and non-neurons (Figure 3e). Further neuronal sub-clustering identified 17 distinct populations, primarily divided into GABAergic (gab1-12) and glutamatergic (glu1-4) neurons, alongside a small population of peptidergic neurons (npep) (Figure 3f/g), consistent with cell-types previously reported in *Mus musculus* hypothalamus^33,34^. Superimposed on these divisions was a complementary organization of cells according to expression of voltage-gated calcium channel subunits (i.e., Cacna2d1, Cacna2d2, Cacna2d3, or none), which frequently emerged as distinguishing marker genes for neuronal sub-clusters (Figure 1g, Supplemental Data File 1, Supplemental Table 9, Supplemental Figure 5). The composition of samples across the five behavioral phenotypes was roughly the same. The relative distribution of nuclei across the 17 neuronal sub-clusters only varied by behavioral phenotype in three pairwise comparisons, and only by 1-3% (Supplemental Data Figure 6).

Given their well-established role in parental behavior, we analyzed the expression of key neuropeptides— including oxytocin, vasopressin, galanin, thyrotropin-releasing hormone—as well as receptors for estrogens, androgens, prolactin, oxytocin, vasopressin, and growth hormone^3,35^. The ‘npep’ cluster was enriched for oxytocin, vasopressin, and growth hormone receptors, while galanin (‘gab6’) and thyrotropin-releasing hormone (‘glu2’) were each enriched in specific neuronal sub-clusters. Several clusters also showed elevated prolactin receptor expression (Supplemental Figure 5). We also examined whether total neuronal expression of these neuropeptides differed across behavioral phenotypes (Figure 3h). Aside from increased Avp expression in allopaternal neurons (driven by a single sample), peptide and receptor expression did not significantly predict individual differences in parental behavior. Thus, MPOA cell-type composition remained stable across phenotypes. These results indicate that natural paternal care was not a result of novel species-specific cell type emergence, and that phenotype was not driven by abundance of particular cell-types. Instead, our subsequent analyses focused on differences in neuronal activity and molecular processes across and within specific neuronal clusters.

### Pup-elicited immediate early gene activity

We next examined whether cluster-specific neuronal activity differed by phenotype to determine whether increased MPOA activity observed among paternal males (Figure 2B) was general or driven by engagement of specific neuronal subtypes. We classified neurons as active based on expression (> 0 counts) of at least one of 9 immediate early genes (IEGs): *Fos, Fosb, Fosl1, Npas4, Egr1, Egr3, Arc, Jun*, and *Junb*. Some IEGs were more highly expressed than others, with the highest expression observed for *Npas4, Fos*, and *Junb* (Figure 3i; Supplemental Data Figure 8). Consistent with our histological study, *Fos* expression was nearly absent in behaviorally naïve controls, confirming that gene expression is induced by pup exposure. Notably, *Egr1* expression was almost exclusive to allopaternal males, whereas *Arc* was most highly expressed in infanticidal males (Figure 3i). Comparing the relative percentage of any-IEG+ neurons within each cluster, we found significant differences in two clusters: gab5 and gab7 (Extended Data Figure 6). In gab5, which is enriched for *Ntng, Pdzrn4*, allopaternal males had more active neurons than sires (+10.2%), controls (+11.3%), and infanticidal males (+8.7%). In gab7, which is enriched for *Unc5d, Cdh18, Cdh23, Zfhx3*, and *Erbb4*, allopaternal males showed +8.4% more active neurons than controls.

To identify phenotype-specific gene expression profiles within different subsets of activated neurons, we performed pseudobulk differential gene expression (DEG) analysis comparing allopaternal and infanticidal males across the three active neuronal subsets with the greatest expression: *Fos*+, *Npas4*+, and *Junb*+. We separated these populations as IEGs are functionally distinct and affect different downstream processes^36^. In *Fos*+ neurons, infanticidal males had significantly greater expression of the gene *Agouti* (agouti signaling protein) (Supplemental Data Figure 8). Within *Npas4*+ neurons, *Agouti* was also the most enriched gene in infanticidal males (followed by *Sst* and four others), whereas *Trh* (thyrotropin releasing hormone) was significantly enriched in allopaternal males (Supplemental Data Figure 8). Finally, within *Junb*+ neurons, infanticidal males showed increased expression of *Tac1*, dopamine receptor *Drd1, Cacna2d3*, and *Lmo7*. These findings suggest that a balance of peptidergic, neuromodulatory, and channel regulation contribute to phenotype-specific MPOA activity.

### *Agouti* is enriched in infanticidal males

To further elucidate gene expression patterns distinguishing opposing parental phenotypes, we analyzed DEGs between infanticidal and allopaternal males at multiple levels of specificity within the neuronal hierarchy, starting broadly with all neurons, then neurotransmitter type (i.e., glutamatergic neurons, GABAergic neurons), and then by specific neuronal cluster. This iterative approach allowed us to detect broad, parsimonious patterns and more fine-grained differences. Despite functional and anatomical differences across neuronal clusters, a small set of DEGs remained remarkably consistent across nearly all clusters, neurotransmitter type, and at the whole-neuron level. Among these, *Agouti* was the most highly differentially expressed gene across all levels of analysis (Figure 3j; Supplemental Data Figure 9a/b). Indeed, *Agouti* was significantly enriched in infanticidal males across 15 of 17 clusters, spanning both glutamatergic and GABAergic populations (Figure 3j/k/l; Supplemental Data Figure 9c-s). Furthermore, *Agouti* expression negatively correlated with pup care before sacrifice (Figure 3m).

To validate this finding, we assessed an independent sample of 23 sexually naïve males and confirmed the negative correlation between pup contact and *Agouti* expression in the MPOA via qPCR (Supplemental Data Figure 10). We also used fluorescent in situ hybridization (i.e., RNAscope Fluorescent Multiplex Assay) to determine the spatial distribution of *Agouti* within the MPOA and confirmed its marked increase in infanticidal males (Figure 3n/o).

### Viral over-expression of *Agouti* elicits infanticide

The *Agouti* gene encodes a paracrine signaling peptide, Agouti signaling protein (ASIP), which is best known for its role regulating pigmentation in vertebrates. *Agouti* is an endogenous antagonist of melanocortin receptors (MCR) with high levels of affinity for MC1R, MC3R, and MC4R37. While ASIP has not been previously linked to infanticidal behavior, its homologue, Agouti-related protein (AgRP), binds to MC3R and MC4R in the brain, increasing cAMP levels and suppressing parental behaviors in Mus dams^38^. This raises the intriguing possibility that *Agouti* may similarly influence environmentally-driven paternal phenotypes in striped mice.

To determine the causal role of *Agouti* in infanticidal behavior, we generated adeno-associated viruses to overexpress the striped mouse *Agouti* cDNA sequence or eGFP alone as a control (AAV2/9-hSYN-AGOUTI-T2A-eGFP or AAV2/9-hSYN-eGFP, respectively) in neurons of the MPOA via bilateral stereotaxic surgery. To ensure consistency with our sequencing study, we focused on GH sexually naïve males, testing the hypothesis that elevated neuronal *Agouti* expression would shift behavioral phenotypes toward infanticidal. Adult sexually naïve males (≥ PND90) reared in GH conditions were pretested and only those displaying ambivalence toward pups were included. This design allowed us to track intra-individual behavioral change from a neutral baseline centered between the two theoretical extremes of caregiving (upper ceiling) and infanticide (lower bound). At both 7-and 21-days post-injection, males were re-exposed to pups, and their behavior was quantified. Successful targeting of the MPOA was confirmed with immunohistochemistry for eGFP, and *Agouti* overexpression was validated by RNAscope using custom probes targeting striped mouse *Agouti* (Figure 4b). *Agouti* overexpression drove a significantly higher incidence of infanticidal behavior compared to both baseline behavior and to the GFP control group (Figure 4c), corroborating predictions from our sequencing analyses. Together, these findings demonstrate that *Agouti* overexpression in MPOA neurons is sufficient to induce infanticidal behaviors in striped mice.

**Fig. 4.**
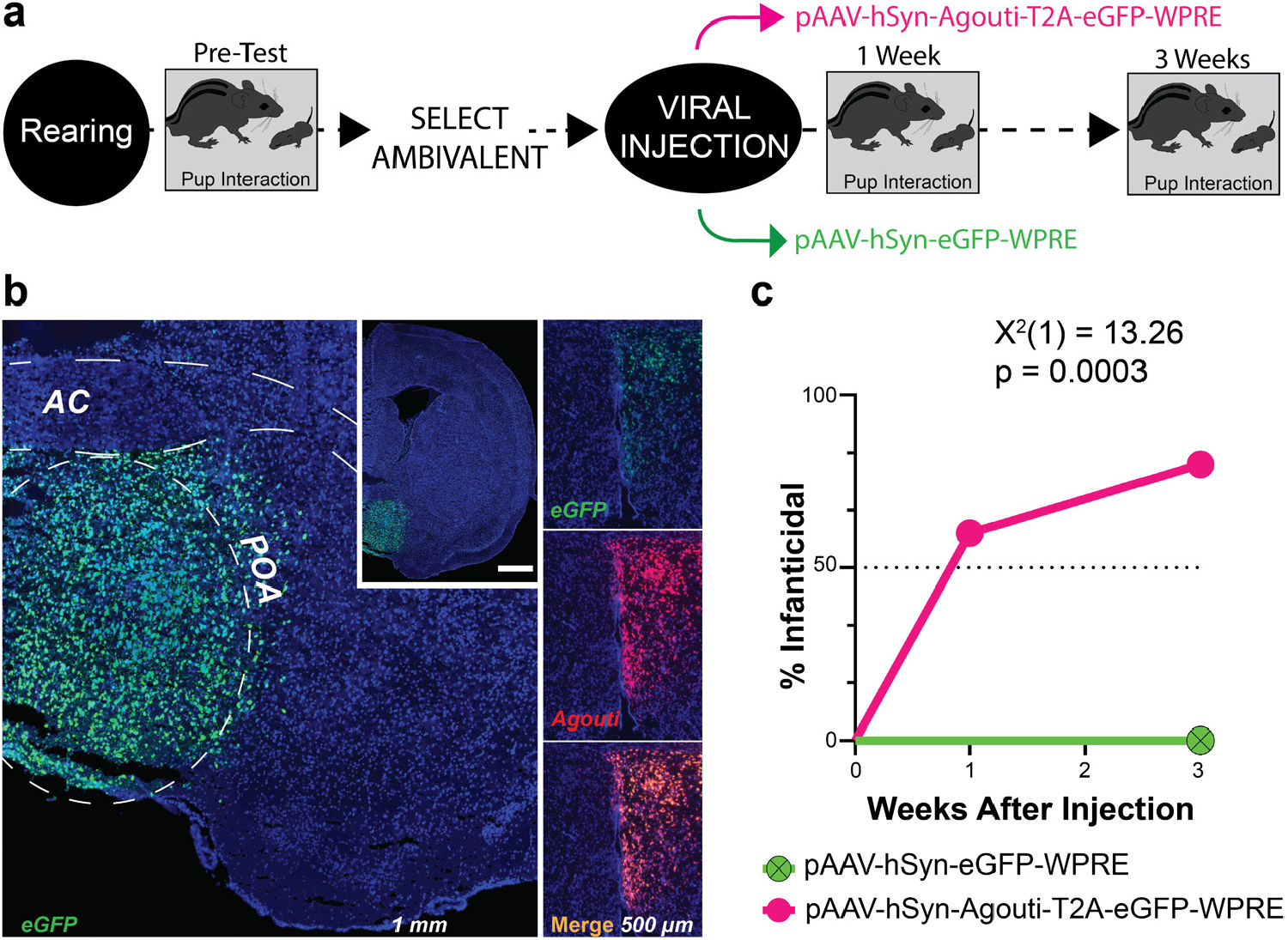
Viral overexpression of agouti elicits infanticidal behavior. (a) Experimental design. Overexpression was confirmed with RNAscope probes for Agouti and eGFP. (b) Example eGFP and Agouti overexpression in a unilateral injection. (c) Proportion of males displaying infanticidal behavior after overexpression of either eGFP or Agouti vectors (Log-rank (Mantel-Cox) test, X2=13.26, df = 1, p= 0.0003). Abbreviations: anterior commissure (AC); preoptic area (POA).

### Agouti integrates socio-environmental information

Ectopic *Agouti* expression in multiple tissues, including hypothalamic brain regions densely populated with melanocortin receptors, induces obesity^39–41^. The ASIP homologue, AgRP, is also linked with the hypothalamic control of feeding. We reasoned that elevated *Agouti* among a subset of infanticidal GH males potentially reflected increased hunger and a motivational state change away from natural paternal behaviors. Indeed, striped mice are maintained on a restricted diet (4g/food/animal/day) to prevent overfeeding and obesity, and GH mice may experience a limited food economy without safeguards against unequal distribution. However, GH males also experienced shared territories compared to SI males, and both food availability and perceived territory are aspects of reproductive fitness that may influence reproductive behaviors including paternal behaviors. To disentangle these two potential drivers of elevated *Agouti* and infanticide, we next manipulated food availability and housing density.

To test the impact of food availability, males were assigned to either a standard diet or a restricted diet (−25%, 3g/animal/day) for 16 days before behavioral testing. To test the impact of housing density, we compared males reared and maintained in GH or SI, as well as those reared in GH but rehoused to SI for the 16 days before testing (GH-SI) (Figure 5a). All animals were tested in a fasted state approximately 24-hours after their last feeding. Before manipulation, SI males remained more parental than GH males, despite fasting, reinforcing our previous findings on rearing environment and paternal care (Supplemental Data Figure 11a-e). At baseline, SI males exhibited significantly lower MPOA *Agouti* expression than GH males (Supplemental Data Figure 11f). Additionally, we compared MPOA *Agouti* levels in GH males sacrificed after fasting (n=6) versus acutely after feeding (≤ 2 hours, n=6) and found no significant difference, indicating that *Agouti* expression is not sensitive to acute changes in satiety (Supplemental Data Figure 11g).

**Fig. 5.**
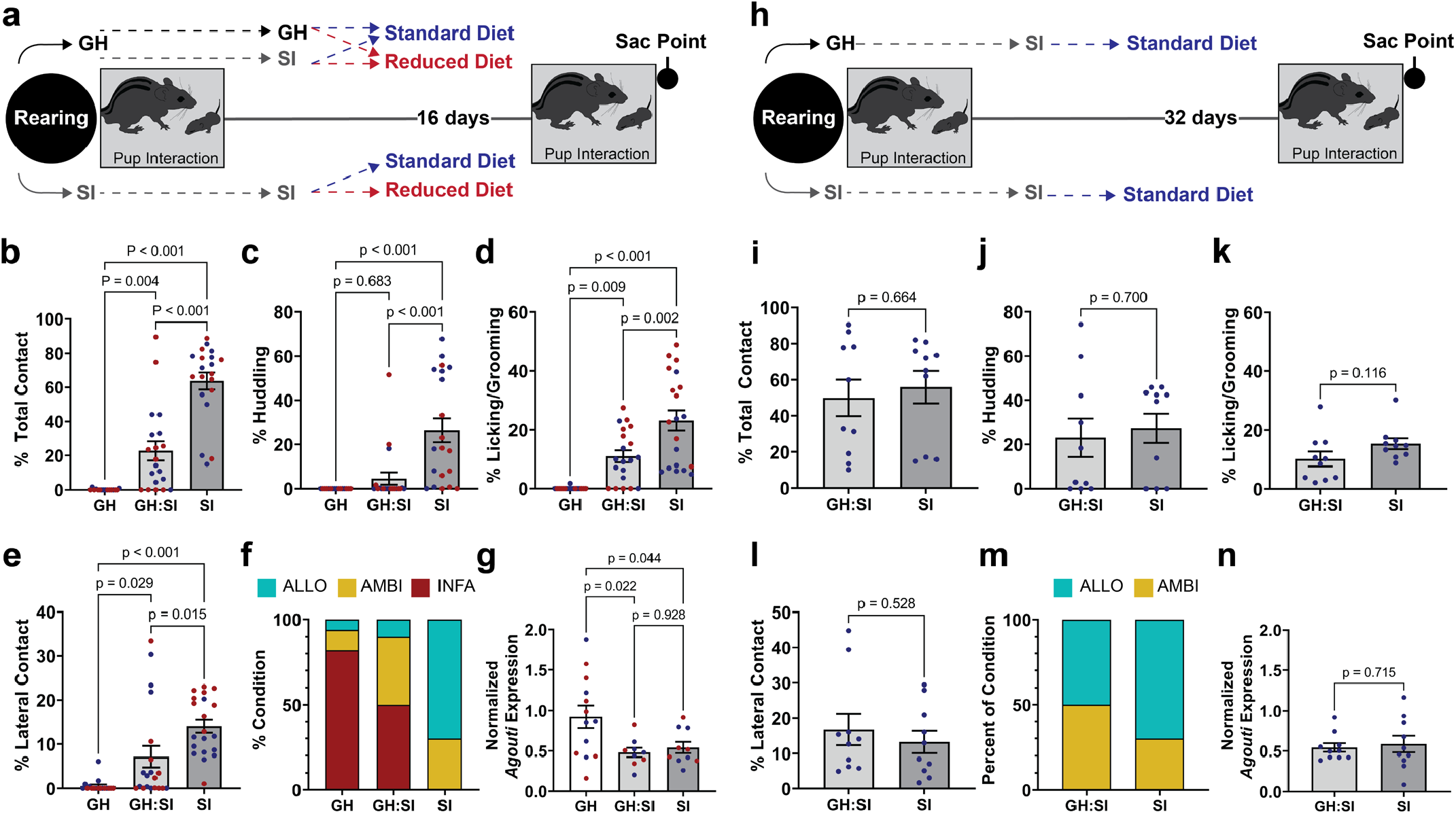
Agouti integrates socio-environmental cues. (a) Schematic of experimental design for feeding and housing manipulations. (b-f) Behavioral outcomes across housing conditions, including (b) total caring contact and constituent behaviors of (c) huddling, (d) licking/grooming, and (e) lateral contact. Animals included SI males (n=20), GH males (n=23), and GH-SI males (n=20). Feeding condition is denoted by color: reduced diet (red); standard diet (blue). Only p-values for pairwise comparisons within the main factor of housing are displayed. (f) The distribution of parental (turquoise), ambivalent (yellow) and infanticidal (red) phenotypes across housing conditions. Comparisons between SI and GH males (X2= 9.643, df = 2, p = .008); and between SI and GH-SI (X2= 18.10, df = 2, p = 0.0001); and between GH-SI and GH males (X2= 2.229, df = 2, p = 0.1883). (g) Expression of Agouti in the MPOA across housing conditions. (h) Schematic of experimental design for prolonged re-housing experiment where a subset of striped mice were maintained for an additional 16 days in SI with standard diet and tested for paternal behaviors (i-m) and MPOA Agouti expression (n). Error bars represent standard error of means.

Following 16 days of manipulation, food-restricted males lost an average of 8.5% of total body mass within 16 days (Supplemental Data Figure 11h), a decline comparable to seasonal weight loss observed during the transition from the wet breeding season to the dry season^42^. Fasted serum levels of leptin, a hormone which positively correlates with satiety^43^, were associated with weight change across the manipulation period, confirming increased hunger (decreased satiety) in food-restricted animals (Supplemental Data Figure 11i). Food restriction did not significantly impact paternal behavior, except for increased licking/ grooming in food-restricted SI males compared to standard-fed SI males (p_adj_ = 0.0005) (Figure 5b-f). Normalized MPOA *Agouti* expression did not differ by feeding condition (p= 0.560) (Figure 5g). This was surprising, given previous associations between *Agouti* and food consumption.

By contrast, recent housing manipulation for 16-days did alter paternal care. Consistent with our previous findings, SI males still displayed the highest levels of total contact, huddling, licking/grooming, and lateral contact, and GH males showing the lowest levels of all parental outcomes. Recently rehoused GH-SI males exhibited intermediate levels of care, increased from the average GH level of care but still lower than average SI levels of care (Figure 5b-e). Similarly, 16 days of single housing reduced, but did not eliminate infanticidal behaviors from GH-reared males (Figure 5f). In addition, normalized MPOA *Agouti* expression was significantly lower in GH-SI males than GH males and comparable to that of SI-reared males, suggesting that *Agouti* was sensitive to the change in housing density (Figure 5g).

Given that the behavior of GH-SI males appeared to shift away from dysparental toward allopaternal behaviors in response single housing, we sought to further assess the temporal effects of our manipulations with an extended rehousing period. A subset of SI and GH-SI males from both feeding conditions was maintained in SI housing and fed a standard diet of 4g/day/animal for an additional 16 days (Figure 5h). This extended period of isolation eliminated prior behavioral differences between SI and GH-SI males, with infanticide absent in all animals (Figure 5i-m). MPOA *Agouti* expression was also not different between SI and GH-SI males (Figure 5n). Taken together, our 16-day and 32-day manipulation experiments support *Agouti*’s role as an integrator of socio-environmental information (e.g., population density or perceived territory) rather than one of food availability or satiety state.

## DISCUSSION

In this study, we leverage the natural paternal behavior of striped mice to investigate the neural and molecular mechanisms underlying naturally occurring paternal care. Through behavioral and histological assays, we demonstrate that MPOA activity distinguishes environmentally driven variation in paternal behaviors, while transcriptomic analyses and causal manipulations revealed that *Agouti* expression in MPOA neurons drives infanticidal behavior. Furthermore, we demonstrate that *Agouti* integrates socio-environmental information to regulate paternal behavior.

Natural variation in allopaternal care in striped mice facilitates the study of male parenting. In contrast to traditional murine models (*Mus musculus* and *Rattus norvegicus*), which require sensitization or reproductive experience to exhibit pup-tolerance and caregiving^8–11^, we show that lab-reared male striped mice display natural paternal care without prior pup exposure, some even at equal levels to sires and dams ^3,18,44^. Our findings align with field studies showing that striped mice adjust parental behavior based on ecological pressures, including population density and resource availability^21,22,24,44^. Our results confirm that socio-environmental factors (GH vs. SI rearing) influence paternal behavior, with social isolation (SI) increasing rather than decreasing paternal behaviors—a stark contrast to the anxiety-like phenotypes observed in *Mus* and *Rattus* following social isolation^45,46^

Our findings support the hypothesis that male parental care in striped mice results from coopted maternal care mechanisms rather than independently evolved ones^3,6,7,15,35,47–52^, ruling out evolution of novel cell types or radically different circuit architecture. Pup exposure elicits cFOS activity within virgin male MPOA as in postpartum murine females, sensitized virgin females, and males of other naturally paternal species [e.g., prairie voles (*Microtus ochrogaster*) and *Peromyscus*]^5,53,54^. Here, we also find that level of MPOA cFOS activity reflects individual differences in striped mouse paternal care, and that correlated activity across brain regions differs between dysparental and allopaternal males.

Within the MPOA, our molecular characterization clearly reveals *Agouti* acts as a transient “off switch” for allopaternal care. As MPOA Agouti levels naturally increase, paternal care decreases and infanticide increases. This causal relationship is further demonstrated by induction of infanticide in previously tolerant males via viral overexpression of *Agouti* within the MPOA. Moreover, although the negative relationship between *Agouti* and paternal care was discovered from sequencing performed within only group-housed males, *Agouti* expression indeed links ecologically relevant socio-environmental conditions with phenotype variation. MPOA *Agouti* is naturally lower among striped mice reared in social isolation, but a transition from group to solitary housing even in adulthood likewise allows *Agouti* levels to fall and paternal care behaviors to reemerge. Notably, this GH-to-SI transition mirrors the behavior of wild male striped mice, who after dispersal from their natal nest form bachelor groups, but abandon these bachelor groups in favor of solitary wandering or territorial breeding^24^. Field evidence shows that bachelor groups form primarily when there is a high male-to-female ratio or when population density is low, and these groups are maintained for 1-5 month periods usually during the breeding season^24^. Thus, situated within this natural ecological context, we speculate that standard laboratory group housing of striped mice emulates natural cues of female sparsity or population density, which in turn elevates MPOA *Agouti* and inhibits allopaternal care. By contrast, the transition into social isolation signals environmental change, reduces MPOA *Agouti*, and lifts inhibition of allopaternal care.

The identification of MPOA *Agouti* as an integrator of socio-environmental information (but not hunger) to suppress paternal care was surprising, particularly because its function in the brain is poorly understood. Agouti-signaling protein (ASIP) is canonically known for its competing role with alpha-MSH at MC1R at the surface of melanocytes within the skin where it prevents the production of eumelanin and stimulates the production of pheomelanin^37^. ASIP is also known for its contributions to obesity in both adipose tissue^40,41,55^ and the brain^39^. Moreover, ASIP’s homolog, Agouti-related protein (AgRP), functions centrally in the arcuate nucleus of the hypothalamus by binding MC3R and MC4R to elevate cAMP and promote feeding^37^. Notably, AgRP+ projections to the MPOA inhibit parental behavior in Mus dams, representing a trade-off between energy expenditure (pup care) and energy acquisition (feeding)^38,56^. Given ASIP’s high affinity for MC1R, MC3R, and MC4R, *Agouti* likely exerts its effects within the brain—either locally in the MPOA or downstream through MC3R and/or MC4R— to regulate parental behavior. However, our findings suggest that, unlike AgRP, *Agouti* is more strongly influenced by socio-environmental cues rather than transient hunger states. Moreover, food-restriction disrupts and reduces dams’ maternal behaviors, mediated by AgRP+ neurons^56^, but does not impinge on striped mouse paternal behavior, suggesting distinct roles and time-scales for AgRP and *Agouti*. Here instead, our findings indicate that long-term environmental conditions influence *Agouti* levels which in turn help MPOA switch between aggression (high *Agouti*) and caregiving (low *Agouti*) states. Given the competing tradeoffs between parental investment and individual survival^57^, *Agouti* may represent an evolutionary mechanism to integrate social information (e.g., population density or social competition) and accordingly shift the behavioral balance between self preservation and offspring investment.

## ACKNOWLEDGEMENTS

This research was supported in part by the Princeton Dean for Research New Ideas in the Natural Sciences (CJP), New York Stem Cell Foundation (CJP), NICHD F32 HD110180 (FDR), Vallee Foundation (RM) and NIH R35GM133758 (RM). CJP is a New York Stem Cell Foundation Robertson Investigator and Robin Chemers Neustein Fellow. We would like to thank Oliver Huang and Angela Chan of the Princeton Neuroscience Viral Core for assistance generating the AAV Agouti and -eGFP over-expression vectors. An initial modification of the nuclei isolation protocol was generated by Adelaide Minerva. We would also like to thank Wei Wang, Jennifer Miller, and the rest of the Genomics Core Facility in the Lewis-Sigler Institute for Integrative Genomics at Princeton where RNA-seq was performed. Thank you to Y. Andre Wang, Karen Bales and the members of her lab, Dr. Zoe Donaldson, and Alexander Ophir and the members of his lab for helpful conversations regarding interpretation throughout the development of this work. Immense thanks is due to the members of the Mallarino and Peña labs, including undergraduates Eman Ali and Daniel Choo, for their support and feedback.

## AUTHOR CONTRIBUTIONS

These studies were designed by FDR, CJP and RM. Data were collected by FDR, CJP, RM, SM, AK, and SK. Data were analyzed by FDR. The manuscript was written by FDR, CJP, and RM with input from all authors.

## METHODS

### Animals

All experiments were conducted in accordance with the guidelines of the Institutional Animal Care and Use Committee at Princeton University and the guidelines and policies of the United States Department of Agriculture, as required by the Animal Welfare Act. Subjects included laboratory-bred African striped mice (Rhabdomys pumilio, or “striped mice”), which originated through systematic outbreeding in a captive colony at Princeton University (obtained via Harvard University, in turn obtained via University of Zurich) and originating from Goegap Nature Reserve, South Africa (S 29° 41.56′, E 18° 1.60′). Striped mice were maintained in a temperature- and humidity-controlled vivarium on a 12:12 light-dark cycle (lights ON at 07:00; lights OFF at 19:00) with water ad libidum. Standard rodent chow (LabDiet 5053, PicoLab Rodent Diet 20) was restricted to 4g per animal per day to prevent the onset of obesity. Sunflower seeds (approximately 10 seeds per animal) were also provided as a form of enrichment. All striped mice were housed in large polycarbonate rat cages (30.80 cm x 59.37 cm x 22.86 cm, Thoren) with a corncob/cotton-mixed bedding with a small volume of shaved aspen and sterile cotton squares for additional nesting material. Additional enrichment in the home cage included a translucent red plastic tube and a plastic running wheel. Sexually naïve striped mice were housed between 1-5 per cage. Breeding animals (including dams and sires) were pair-housed with mates and pups under the conditions described above. Sex was assigned as male or female according to visual inspection of externalized genitalia and anogenital distance at weaning on postnatal day (PND) 20 and confirmed in adulthood (≥ PND 90) with the presence or absence of testes.

### Statistical Software and Power Analysis

#### Statistical Sofware

Analyses were completed in R49 (version 4.4.1) and Graphpad Prism (version 9.4.1). For all inferential statistics, a threshold of p ≤ .05 was used to determine significance. Additional methods regarding statistical analysis can be found in supplementary information.

#### Power Analysis

Power analysis for initial paternal behavior testing used the pwr.anova.test function (R, pwr package). Initial parameters (power = 0.80, Cohen’s f = 0.585) were based on published paternal behavior studies19 and calculated to need n=12.5. Parameters were then adjusted based on our own first cohort (n=47: 24 SI, 23 GH, new Cohen’s f=0.606, target power = 0.80) and calculated to need n=11.7 animals per group in a two-way comparison and a 9.8 animals per group in a three-way comparison.

### Group Housing (GH) and Social Isolation (SI)

All animals were reared with both biological parents until weaning. At weaning, male offspring were assigned to either “group housed” (GH) or “socially isolated” (SI) conditions. GH males were weaned into new cages in groups of 3-4 similarly aged conspecifics (+/-3 days). SI males were weaned into a new cage and housed alone. All other housing conditions (e.g., enrichment, feeding, etc.) were equal and maintained as described above. No more than one littermate was used per condition in each experiment to experimentally control for litter effect.

### Behavior

Sexually naïve male striped mice underwent open field and social interaction behavioral testing from PND80-90. Pup interactions were assayed at ≥PND90 (i.e., gonadal maturity).

#### Open Field Test

Open field testing (OFT) was used to quantify exploratory and anxiety-like behaviors 59,60. Striped mice were allowed to explore an open 44×44 cm arena for 10 minutes, as previously described60. Because African striped mice are diurnal, testing occurred under light conditions between 12:00 and 17:00. Total distance traveled and time spent in the center (24 cm x 24 cm) of the arena were recorded by top-down camera and measured via Ethovision software (Noldus).

#### Social Interaction Test

A social interaction test was adapted from standard protocols as previously described60. Here, striped mice explored a 44×44 cm arena for two 150-second trials. In the first, an empty wire-mesh enclosure was centered on one wall; in the second trial an adult male conspecific was placed into the enclosure. Two males (both littermates of similar size) were used as stimuli for all test subjects. Time spent in the “interaction zone” (14x 26 cm) surrounding the enclosure and “distance traveled” within the arena were recorded and quantified using Ethovision software (Noldus). A social interaction ratio (SI Ratio) was calculated as time spent exploring the novel mouse over time exploring the empty enclosure. All social interaction testing was conducted in the light phase between 12:00 and 17:00.

#### Pup Interaction Test

Parental care was assayed with a 20-minute pup interaction test, which was adapted from the alloparental care test used in prairie voles61. Testing occurred in a large rat cage (30.80 cm x 59.37 cm x 22.86 cm, Thoren) with a thin layer of cotton square bedding. Animals were habituated to the cage for 40-minutes. Following habituation, the test animal was relocated to one end of the cage and maintained within a large cup (≤ 5 seconds) while a novel pup stimulus (PND1-6) was placed in at the other extreme of the cage. Each tested male was released to interact freely with the pup. If the test animal attempted infanticide, test was immediately ended and the pup either returned to its parents (if unharmed) or euthanized via emergency decapitation. Notably, wild striped mice do not appear to distinguish unfamiliar pups from their own before PND10^62^. We did not control for pup sex, but we selected pups from their home nest at random. Based on observations, neither male nor female pup stimuli are more likely to be attacked by sexually naïve males (X2=0.06857, p=0.7934, from sample of 60 tests of adult, sexually naïve males with 30 male and 30 female pup stimuli). Specific outcomes included durations and latency to the first presentation of key behaviors: inspection, licking/grooming, lateral contact, huddling, attack (i.e., infanticide), and total caring contact (huddling + lateral contact + licking/grooming). Pup interaction tests were completed in the light cycle. Animals were fed prior to testing with the exception with those included in the experimental manipulation of food availability. All behavior was digitally recorded using a top-down digital webcam (Logitech BRIO, ASIN: B01N5UOYC4). Behavior was manually scored by a trained observer using the Behavior Tracker software (http://behaviortracker.com/), provided courtesy of Dr. Karen Bales.

#### Initial Pup Interaction Testing, Characterization

Initial behavioral characterizations of parental care were made with a sample of sexually naïve male striped mice conditioned as GH (N = 24) and SI (N = 23), as well as primiparous striped mouse fathers, i.e. sires (N = 18), and mothers, i.e. dams (N = 7).

#### Categorization of behavioral phenotypes

Sexually naïve males were phenotypically categorized as infanticidal, ambivalent, or allopaternal. Males were categorized as infanticidal if they demonstrated pup-directed, aggressive behaviors requiring researcher intervention. Allopaternal was defined as spending ≥ 10% of the test duration in any caring contact and ≥ 5% of the test duration in huddling contact. Ambivalent males were those that were neither infanticidal nor Allopaternal.

#### Statistical Analysis

Analyses for OFT and social interaction were completed in R (version 4.4.1). Mixed effects models with a random effect for litter were used to analyze all duration-based measures to account for litter effects where necessary. Analyses for pup interaction tests were completed in Graphpad Prism (version 9.4.1). Only one male per litter per experimental condition were used for pup interaction tests. Latency data were analyzed with survival curve analysis (Log-rank/Mantel-Cox test). Duration data were assessed with unpaired two-sided t-tests or ANOVA. Compositional differences (i.e., for phenotypic category) were determined by chi-square difference test.

### Naturalistic Intervention Study

Adult sexually naïve GH-or SI-reared males were rehoused into social isolation (n=20 each); a second group of GH males were maintained in group housed conditions (n=24). Half of each group were provided a standard diet of 4g food per day per animal, while the other half received 3g/ day/animal. Animals were maintained in these conditions for 16 days with weights taken every 3 days. Each animal was tested on OFT and pup interaction tests before and after rehousing/food restriction. Brain and blood tissue was collected from half the animals immediately after the second pup interaction test. The remaining animals were all maintained under a standard diet (4g/day) in their housing conditions from the previous 16 days. These animals were behaviorally tested a third time prior to tissue collection.

### Brain-wide cFOS expression mapping

#### Tissue Collection

Between PND90 and PND95, 22 males (n=11 GH, n=11 SI) completed a second pup interaction task in which they were continuously exposed to the pup for 40-minutes before sacrifice. If males attempted infanticide, a new pup was placed in a tea strainer within the testing cage to allow for continued exposure. 20 additional males (n=10 GH, n=10 SI) were used as controls and were exposed to an empty cage. At the end of the 60-minute exposure or control period, animals were anesthetized via IP injection of ketamine (100mg/kg) and xylazine (20mg/kg) and transcardially perfused with 1mL/g 1XPBS followed by 1mL/g ice-cold 4% paraformaldehyde. Brains were extracted and stored in 4% paraformaldehyde at 4ºC overnight. Brains were transferred to a series of incubations at 4ºC in 10%, 20%, and 30% sucrose before flash freezing on dry ice in O.C.T. compound within a plastic mold and stored at -80ºC. Prior to tissue dissection, brains were habituated to -20ºC overnight in a -20ºC freezer. Tissue was then sliced in sequential, 30µm coronal slices in 120µm intervals and directly mounted to Superfrost Plus microscope slides. Tissue was subsequently stored at -20ºC for short-term storage before immunohistochemistry.

#### Immunohistochemistry

Tissue was removed from -20ºC storage and washed three times in 1XPBS. Tissue was then incubated at room temperature for 1-hour with 1.5% normal donkey serum (NDS) in 0.25% Triton-X100 in 1X PBS (PBT). Subsequently, tissue was incubated overnight at 4ºC with primary antibodies diluted 1:2000 in 1.5% NDS in PBT. Primary antibodies were against cFOS (Cell Signaling, Rabbit mAb 2250S) for all slides, and Tyrosine Hydroxylase (ImmunoStar, mAb Mouse 22941) for posterior sections to help in the identification of the ventral tegmental area (VTA). The following morning, the tissue was washed three times in PBT before incubation for 1-hour at room temperature with secondary antibodies were diluted 1:500 in PBT (Jackson ImmunoResearch 711-546-152 and 715-175-150). Tissue was then washed two times with PBT, counterstained with DAPI (0.1%) in PBT for 15-minutes, and then washed twice more in PBT before subsequent drying and coverslip (#1.5) application with Fluoromount-G mounting medium.

#### Imaging and Quantification

Imaging was performed with a slide scanner (Hamamatsu NanoZoomer S60, C13210-01) using a 40X objective lens and filters or DAPI, FITC (cFOS), and Cy5 (Tyrosine hydroxylase, for posterior sections) but exported at 10X for subsequent processing using FIJI63. All brain slices were assigned to corresponding Paxinos mouse brain atlas sections64. For each region of interest (ROI), researchers quantified bilateral densities of cFOS+ neurons in a semi-automatic process. Researchers manually traced ROI outlines and recorded ROI areas (mm2). Images were thresholded using a FIJI macro before cell count quantification with a second FIJI macro. Density was calculated as cFOS+ neurons / area for each ROI.

#### Analysis

CFOS+ density was analyzed by 2×2 ANOVA with main effects and interactions for housing (GH vs. SI) and pup-exposure (exposed vs. not exposed), or by 1-way ANOVA with a main factor of phenotype (parental, dysparental, or control). Post-hoc, pairwise comparisons were only performed given significant main effects or interactions (p<0.05). Outliers were determined at a sample level (agnostic of group or phenotype) removed region-by-region prior to ANOVAs. For correlation analysis, cFOS+ density for pup-exposed animals was standardized to mean control cFOS+ density for each ROI separately. Because the distribution of the density data was not normally distributed and because our questions were related to rank order, Spearman’s rank-order correlation was calculated.

### Single Nucleus RNA-Sequencing

#### Collection of Tissue

MPOA of 17 striped mice were processed for single nucleus RNA-sequencing (snRNA-seq): Sexually naïve adult males that were infanticidal (n=3), allopaternal (n=3), or ambivalent (n=3), as well as dams (n=4) and their male mates (n=4, ‘sires’). Following a baseline pup interaction test, all dams, sires, infanticidal, and allopaternal individuals were exposed again to a pup stimulus for five minutes following onset of interaction (variable time to initiate interaction). Dams and sires were exposed to their own offspring, while infanticidal and allopaternal males were exposed to novel pup stimuli. Because the onset of infanticide was rapid, infanticidal males were continuously exposed to a second, novel pup stimulus held in a tea strainer from the time of infanticide until the end of the 5-minute exposure period. Following the exposure, animals were euthanized by rapid decapitation. Rather than experiencing a second pup exposure, ambivalent males were instead sacrificed from an empty cage. Brain tissue was extracted and rinsed in ultra-pure 1XPBS before sectioning into 1mm tranches in an ice-cold brain matrix. The MPOA was bilaterally dissected with a 14-guage punch and flash frozen in RNase-/DNase-clean 1.5mL tubes on dry ice. Tissue was collected and frozen within 2-mintues of sacrifice. All tissue punches were stored at -80ºC until subsequent processing.

#### Isolation of nuclei, library preparation, and sequencing

Nuclei were isolated from frozen tissue punches in batches of 4 samples, counterbalanced by phenotype. The protocol for nuclei preparation was modified from 10X protocol GC000393. Samples were Dounce homogenized 10 times in 500µl of homogenization buffer with a 2ml Kimble Kontes Dounce tissue grinder on ice. 5% IGEPAL CA-630 was added to the Dounce and each sample was further homogenized 5 times before filtering through a 40µm cell strainer (BD Falcon 352340). The filtrate was then diluted 1:1 with a 50% Iodixanol solution (OptiPrep Density Gradient Medium (Sigma D1556) diluted to 50%) and added to a 1.5mL microcentrifuge tube containing 100µl layers (from bottom to top) of 40% and 30% Iodixanol. Samples were then centrifuged at 4500rcf in a swinging bucket centrifuge at 4ºC for 45-minutes with a moderated ramp-up speed and the lowest break setting. Nuclei were then extracted in a 60µL volume from the resulting interstitial layer between the 40% and 30% Iodixanol layers. Nuclei concentrations were then determined via hemocytometer. Isolated nuclei were transferred to the Genomics Core Facility in the Lewis-Sigler Institute for Integrative Genomics at Princeton for library preparation with the 10X Genomics Chromium system and sequencing. Libraries were prepared based on an estimated input of 20,000 and recovery of ∼12,535 nuclei per sample. All subsequent sequencing was carried out on an Illumina NovaSeq 6000 and samples were sequenced to a mean depth of 42,426 reads per nucleus (SD=8894) and a mean saturation of 71.64 (SD=10.01).

#### Data Analysis

Data was demultiplexed by barcode and processed with 10x Genomics Cell Ranger, CellBender65 and the R package scDblFinder^66^ to filter and remove doublets and nuclei with high mitochondrial reads or ambient RNA and random barcode swapping. Subsequent bioinformatic steps were carried out in the R package Seurat^67^. Seurat objects were created for each original identity (i.e., biological sample). There were 164,507 nuclei that passed filtering and were included in the final dataset. There was a mean 1985 UMI/nuclei and 1206 genes/nuclei.

Prior to sample integration, cells were renamed to add cell identifiers for each biological sample to account for overlapping barcodes across different biological samples. Individual samples were then merged, normalized, scaled, and integrated using the harmony 32 package. This integrated dataset was then iteratively clustered within a range of dimensional embeddings initially determined through visual inspection and identification of a plateau within an ElowPlot of the standard deviation of the first 50 principal components. Subsequently resolution was adjusted until the resulting cluster boundaries were congruent with the expression of marker genes, as visualized with the feature plot function in Seurat. Accordingly, cells were initially separated into 8 major cell classes, including astrocytes, ependymal, endothelia, microglia, myelinated oligodendrocytes, new oligodendrocytes, oligodendrocyte precursors, and neurons. Separation of cell clusters was accomplished using Seurat clustering and examination of enriched expression of known class marker genes (e.g., *Aqp4, Tmem212, Cldn5, Cx3cr1, Mog, Pdgfra, Enpp6*, and *Rbfox3*, respectively).

Neurons were separated and the neuron-specific data was re-clustered and the resulting UMAP was visually examined. One neuronal population was identified as rare, with presence in only a small subset of individual across behavioral phenotypes, showing incongruency within phenotype and across the broader sample. Another population of neurons were highly congruent and showed obvious enrichment for genes associated with cholinergic neurons (e.g., Chat). To reduce variance and improve delineation between GABAergic and glutamatergic neurons, this and the rare cluster were removed and the integrated dataset was clustered again. The resulting, filtered dataset was then re-clustered with 45 dimensions and a resolution of 0.35 with the original Louvain algorithm. This resulted in a final clustered dataset with 26 neuronal clusters. The delineation of these clusters was initially considered through a consideration of cluster-specific gene enrichment via the FindAllMarkers function with the ROC algorithm in Seurat with a logFC threshold of 0.1 and a requisite of a minimum of 10% of cells in a cluster expressing an enriched gene. This initial assessment demonstrated that several clusters lacked significantly distinguishing features from neighbors; thus, those poorly distinguished clusters were merged. The resulting neuronal clusters demonstrated cluster-specific enrichment (Supplemental Table 2, Supplemental Data File 1) and were primarily delineated by neurotransmitter type: i.e., GABAergic (enriched for *Slc32a1, Gad1, Gad2*), Glutamatergic (enriched for *Slc17a6*), or Neuropeptidergic (enriched for *Avp, Oxt*, and *Caprin2*). A one-way ANOVA was run for each cluster to determine if there were differences at the level of phenotype in cluster composition. If a significant effect were found at the level of the omnibus test, post hoc pairwise comparisons with Tukey HSD correction were made to identify the source of variance between groups. Pseudobulk analysis was carried out to address questions about molecular processes within neurons more generally (and within smaller subsets) and in active neuronal populations. To the former, three subsets were created by selecting for all neuronal clusters or for clusters specifically labeled as GABAergic (e.g., gab1… gab_n_) or glutamatergic (e.g., glu1 … glu_n_). To the latter, subsets were created according to immediate early gene (IEG) expression (> 0 counts recorded of each respective IEG in a given nucleus). IEGs were considered separately, as the cellular processes associated with each IEG are distinct^36^. Those IEGs considered included the following: *Arc, Egr1, Egr3, Fos, Fosb, Fosl1, Jun, Junb*, and *Npas4*. The primary contrast for pseudo bulk analysis was that of alloparental and infanticidal sexually naïve males. Pseudocounts were extracted using the AggregateExpression function in Seurat with subsequent application of the DESeq2^68^ modification of the function FindMarkers. Pseudocounts were also correlated with continuous behavioral outcomes and correlated with a Pearson’s product moment correlation. Notably, in our consideration of IEGs, pseudobulk contrasts could not be drawn between allopaternal males and infanticidal males for *Egr1* because there were an insufficient number of *Egr1*+ neurons in the infanticidal condition. Likewise, pseudobulk contrasts could not be drawn between allopaternal males and formerly ambivalent, unexposed control males for Fos and *Egr1* because there were an insufficient number of Fos+ and *Egr1*+ neurons in the control condition. Contrasts were drawn neither for *Egr3* nor *Arc*, as there were insufficient numbers of *Egr3*+ and *Arc*+ neurons in allopaternal males.

### Fluorescent In Situ Hybridization

Tissue was harvested as described above for immunohistochemistry, except animals were only perfused with 1XPBS before dissecting brains, and brains were flash-frozen in isopropanol and frozen at -80ºC in blocks of OCT. Tissue was sliced in sequential, 16µm coronal slices and immediately mounted to Superfrost Plus microscope slides and left tissue-side up at -20ºC for two-hours to improve tissue adhesion. Slides were subsequently stored at -80ºC in small slide boxes cleaned a priori with RNase Away, with a small desiccant packet, and sealed with electrical tape until processing. Gene expression was detected by RNAscope Multiplex Fluorescent ISH V2. Probes for Agouti and eGFP were custom made and are commercially available via ACDBio (RNAscope™ Probe-Rpu-Asip-C3, catalogue 1328331-C3; eGFP: ACDBio Catalog 53885*). Tissue was processed according to kit instructions with the following exceptions, excepting that tissue was treated with 10% Protease IV in lieu of 100% Protease IV. Images were initially captured and assessed using a slide scanner (Hamamatsu NanoZoomer S60, C13210-01) using a 40X objective lens and filters or DAPI, FITC, TRTC, and Cy5. Images were subsequently captured at 40X using a Nikon A1R-STED Confocal Microscope with Simulated Emission Depletion Detector.

### qPCR

MPOA tissue punches were collected identically as described above for snRNAseq from sexually naïve GH males (n=23). RNA was extracted from tissue samples with Trizol (Invitrogen) and chloroform (Sigma) and purified using an RNeasyMicro Kit (Qiagen 74004). cDNA was created with a reverse transcription kit (HighCapRT, Applied Biosystems). All real-time semi-quantitative qPCR reactions were run in triplicate and used SYBR-green on a QuantStudio 7 (ThermoFisher). Relative gene expression for striped mouse Agouti was quantified by qPCR and normalized to striped mouse Gapdh using the 2-ΔΔCt method. The primer sequences used for agouti were as follows: Forward: 5’-GGA GAC ACT TGG AGA TGA CAG G – 3’; Reverse: 5’-CTC TTC TGC TTC TCG GCT TC – 3’; And the sequences used for Gapdh were as follows: Forward: 5’ – AGG TCG GTG TGA ACG GAT TTG – 3’; Reverse: 5’ – TGT AGA CCA TGT AGT TGA GGT CA – 3’.

### ELISA

Trunk blood was collected into protein LoBind tubes and allowed to sit undisturbed at room temperature ≥ 30 minutes. Blood was then centrifuged at 1500g for 10-minutes at 4ºC, and serum was subsequently removed and frozen at -80ºC until later use. Serum was processed via ELISA for leptin (Invitrogen KMC2281) according to the manufacturer’s protocol. Results were read on a Tecan Spark plate reader at 450nm (outcome), 540nm (background), and 570nm (background). Before analysis, results of the 540nm reads were subtracted from the 450nm reads. Lines of best fit / standard curves were fit from standards in excel and concentrations were inferred. The mean intra-assay CV was 2.0%. The inter-assay CV was 6.17%.

### Viral Manipulation Study

#### Viruses

All viruses were generated and packaged by the Princeton Neuroscience Institute Viral Core Facility at Princeton University. For Agouti overexpression, we generated AAVs (serotype 2/9), carrying pAAV-hSyn-Agouti-T2A-eGFP-WPRE (titer: 1.8e14 genome copies/ml). As a control, we generated nearly identical vectors lacking the *Agouti* transgene (pAAV-hSyn-eGFP-WPRE; 2e14 genome copies/ml). An hSyn promoter was selected to confer neuronal specificity. Efficacy of the viral transfection was confirmed with RNAscope with probes designed for Agouti and eGFP (see above).

#### Stereotaxic Surgical Procedures

Adult sexually naïve GH males were tested a priori in a pup interaction test. Only ambivalent animals were selected for viral manipulation and were randomly assigned to either Agouti overexpression or control GFP groups. AAVs were infused into MPOA bilaterally via stereotactic surgery. Anesthesia was initiated with an IP injection of ketamine (100mg/kg) and xylazine (10mg/kg) diluted with sterile saline. After 10-minutes, animals were fitted into a dual-arm stereotax with ear bars and maintained on 0.5%-1.5% isoflurane in oxygen via nose cone for the duration of the surgery. To target MPOA, two Hamilton syringes were angled inward at 12º, centered at Bregma, and lowered to AP +1.35mm, ML ± 1.42mm, DV -5.8mm. 150nL of virus was infused per side and allowed to absorb into tissue for 10 minutes before slowly removing syringes. Following surgery, animals were given a single, subdermal dose of Ketoprofen (10mg/kg), and bupivacaine was topically applied to the incision site. Bupivacaine was applied again 24-hours post-surgery. Animals were provided with a single pack of Diet Recovery Gel (ClearH20, SKU: 72-06-5022). Recovery was monitored for five days post-surgery. Pup interaction tests were conducted one and three-weeks post-injection, after which animals were euthanized, and brain tissue was fixed via transcardial perfusion. Viral injection site locations were confirmed with IHC for the eGFP tag (primary Ab: Aves Labs GFP-1020; secondary Ab: JacksonImmuno Research 703-545-155). Animals for whom Agouti overexpression was off-target were included in the control group. Initial efficacy of viral overexpression was determined with RNAscope, as detailed above.

