## Supplemental Material for "*Agouti* integrates environmental information to regulate natural variation in paternal behavior"

**<u>Supplemental Data</u>**

***Content***

**Supplemental Data Tables ............................................................................ 2 - 10**

**Supplemental Data Figures ...................................................................... 11 - 21**

**Supplemental Table 1| Ethogram of Pup Interaction Test Behaviors.**

| Behavior | Description | Type |
|---|---|---|
| Sniff | Olfactory inspection of pup stimulus. Nose in contact with or immediately over pup stimulus at any point of the pups body. | Duration |
| Lick/Groom | Act of licking or grooming the pup, not specific to location on the pup's body. | Duration |
| Lateral Contact | Physical contact with the pup such that the center-point of the pup's mass is exposed. | Duration |
| Huddle | Physical contact with the pup such that the center-point of the up's mass is covered by the body of the test animal. | Duration |
| Infanticide | Physical aggression toward the pup, such that the termination of the test would be necessitated for the survival or execution of a humane endpoint for the pup. | Frequency |

**Supplemental Data Table 2 | Differences in care behavior by phenotype**

| | | Select pairwise comparisons (Tukey HSD) | | | |
|---|---|---|---|---|---|
| **Behavior** | **ANOVA** | **ALLO-INFA** | **ALLO-AMBI** | **ALLO-SIRE** | **ALLO-DAM** |
| **Licking/Grooming** | $F(4,67) = 4.1869$, $p = 0.004$, $\eta_P^2 = 0.20$ | $p = 0.0546$ | $p = 0.138$ | $p = 0.006$ | $p = 0.990$ |
| **Lateral Contact** | $F(4,67) = 9.9239$, $p = 2.29e\text{-}06$, $\eta_P^2 = 0.37$ | $p = 0.00358$ | $p = 0.013$ | $p = 0.889$ | $p = 0.091$ |
| **Huddling** | $F(4,67) = 13.219$, $p = 5.422e\text{-}08$, $\eta_P^2 = 0.44$ | $p = 0.00339$ | $p = 3.36e\text{-}05$ | $p = 0.921$ | $p = 0.046$ |
| **Total Caring Contact** | $F(4,67) = 16.369$, $p = 2.12e\text{-}09$, $\eta_P^2 = 0.49$ | $p = 3.3e\text{-}06$ | $p = 2e\text{-}07$ | $p = 0.782$ | $p = 0.550$ |

Abbreviations: infanticidal (INFA), ambivalent (AMBI), allopaternal (ALLO), genetic fathers (SIRE), genetic mothers (DAM).

**Supplemental Data Table 3 | FOS+ Neuronal Density by region, pup exposure, and housing**

| ROI | Main Effect: Pup Exposure | Main Effect: Housing | Interaction Effect |
|---|---|---|---|
| AON | F(1, 34) = 6.5318, p = 0.01524, $\eta_p^2$ = 0.34 | F(1, 34) = 0.8795, p = 0.35496, $\eta_p^2$ = 0.02 | F(1,34) = 0.3376, p = 0.56504, $\eta_p^2$ = 9.83e-03 |
| BLA | F(1,24) = 1.8103, p = 0.19105, $\eta_p^2$ = 0.28 | F(1,24) = 0.0012, p = 0.97215, $\eta_p^2$ = 0.04 | F(1,24) = 0.6978, p = 0.41178, $\eta_p^2$ = 0.03 |
| CP | F(1, 33) = 0.3141, p = 0.57895, $\eta_p^2$ = 1.93e-03 | F(1, 33) = 4.0382, p = 0.05272, $\eta_p^2$ = 0.09 | F(1, 33) = 1.1275, p = 0.29601, $\eta_p^2$ = 0.03 |
| LS | F(1,34) = 0.0576, p = 0.8118 , $\eta_p^2$= 3.66e-03 | F(1,34) = 1.2509 p = 0.2712, $\eta_p^2$= 0.02 | F(1,34) = 0.4802, p = 0.4931 , $\eta_p^2$= 0.01 |
| LSd | F(1,34) = 0.0660, p = 0.798862 , $\eta_p^2$= 0.01 | F(1,34) = 0.3219, p = 0.574202, $\eta_p^2$= 6.43e-03 | F(1,34) = 0.1137, p = 0.738046, $\eta_p^2$= 3.33e-03 |
| MPOA | F(1,31) = 4.3280, p = 0.04585 , $\eta_p^2$= 0.29 | F(1,31) = 0.0110 , p = 0.08385, $\eta_p^2$= 0.03 | F(1,31) = 0.6878, p = 0.41327 $\eta_p^2$= 0.02 |
| NAcc | F(1,28) = 1.0111, p = 0.32326 , $\eta_p^2$= 0.05 | F(1,28) = 0.2271, p = 0.63740 , $\eta_p^2$= 0.03 | F(1,28) = 0.0293, p = 0.86523 , $\eta_p^2$= 1.05e-03 |
| NAcsh | F(1,29) = 1.2523, p = 0.2723 , $\eta_p^2$= 0.05 | F(1,29) = 0.1907, p = 0.6656, $\eta_p^2$= 7.75e-04 | F(1,29) = 0.1239, p = 0.7274 , $\eta_p^2$= 4.26e-03 |
| PAG | F(1,30) = 0.6210, p = 0.4369, $\eta_p^2$= 0.03 | F(1,30) = 0.2003, p = 0.6577, $\eta_p^2$= 8.25e-03 | F(1,30) = 0.0190, p = 0.8914, $\eta_p^2$= 6.32e-04 |
| PFC | F(1,37) = 12.9154, p = 0.0009445, $\eta_p^2$= 0.49 | F(1,37) = 0.0882, p = 0.7680842 , $\eta_p^2$= 1.58e-03 | F(1,37) = 0.6005, p = 0.4433057, $\eta_p^2$= 0.02 |
| PVT | F(1,27) = 1.2700, p = 0.2697, $\eta_p^2$= 0.14 | F(1,27) = 1.2598, p = 0.2716, $\eta_p^2$= 0.01 | F(1,27) = 0.5702, p = 0.4567 , $\eta_p^2$= 0.02 |
| vBST | F(1,38) = 0.0000, p = 0.9962002 , $\eta_p^2$= 2.95e-03 | F(1,38) = 0.5595, p = 0.4590561 , $\eta_p^2$= 0.01 | F(1,38) = 0.1169, p = 0.7342970 , $\eta_p^2$= 3.07e-03 |
| VP | F(1,35) = 0.2420, p = 0.6258597 , $\eta_p^2$= 7.65e-03 | F(1,35) = 0.6816, p = 0.4146301 , $\eta_p^2$= 0.03 | F(1,35) = 0.0297, p = 0.8641300, $\eta_p^2$= 8.48e-04 |
| VTA | F(1,29) = 3.6804, p = 0.06495 , $\eta_p^2$= 0.26 | F(1,29) = 0.0458, p = 0.83207 , $\eta_p^2$= 2.32e-06 | F(1,30) = 0.1422, p = 0.70888, $\eta_p^2$= 4.88e-03 |

**Supplemental Data Table 4 | FOS+ Neuronal Density by ROI and by pup exposure**

| ROI | Mean (No Exposure) | Mean (Pup Exposure) | T-test | P-value (fdr adjusted) |
|---|---|---|---|---|
| AON | 300.07 | 575.9997 | t(34) = -4.195 | 0.0002 |
| MPOA | 55.32871 | 168.1614 | t(31) = -3.612 | 0.0011 |
| PFC | 160.4113 | 520.6997 | t(37) = -5.929 | < .0001 |

**Supplemental Data Table 5 | FOS+ Neuronal Density by ROI by phenotype**

| ROI | F(DF) | P-value | $\eta_p^2$ |
|---|---|---|---|
| AON | F(2,35) = 8.849 | 0.0007759 | 0.34 |
| BLA | F(2,25) = 5.1391 | 0.013503 | 0.29 |
| CP | F(2,34) = 0.2294 | 0.7962 | 0.01 |
| LS | F(2,35) = 0.1009 | 0.9043 | 5.73e-03 |
| LSd | F(2,35) = 0.2676 | 0.7668 | 0.02 |
| MPOA | F(2,32) = 11.1892 | 0.0002068 | 0.41 |
| NAcc | F(2,29) = 0.9963 | 0.38153 | 0.06 |
| NAcsh | F(2,30) = 1.0186 | 0.373241 | 0.06 |
| PAG | F(2,31) = 1.3128 | 0.2836 | 0.08 |
| PFC | F(2,38) = 18.378 | 2.609e-06 | 0.49 |
| PVT | F(2,28) = 2.3109 | 0.1178 | 0.14 |
| vBST | F(2,39) = 0.234 | 0.7925 | 0.01 |
| VP | F(2,36) = 0.1507 | 0.8606 | 8.30e-03 |
| VTA | F(2,30) = 5.5943 | 0.008613 | 0.27 |

### Supplemental Data Table 6 | FOS+ Neuronal Density by ROI and by phenotype

| ROI | Mean (Control) | Mean (Dysparental) | Mean (Allopaternal) | Control vs. Dysarental | Control vs. Allopaternal | Allopaternal vs. Dysparental |
|---|---|---|---|---|---|---|
| AON | 160.4113 | 544.9152 | 478.3228 | t(35) = -3.859, p = 0.0014 | t(35) = -3.062, p = 0.0063 | t(35) = 0.195, p = 0.8469 |
| BLA | 96.3052 | 177.3848 | 233.5155 | t(25) = -2.061, p = 0.0748 | t(25) = -3.183, p = 0.0116 | t(25) = -1.427, p = 0.1661 |
| MPOA | 55.32871 | 130.7142 | 229.013 | t(32) = -2.353, p = 0.0249 | t(32) = -4.712, p = 0.0001 | t(32) = -2.630, p = 0.0195 |
| PFC | 160.4113 | 544.9152 | 478.3228 | t(38) = -5.705, p < 0.0001 | t(38) = -3.942, p = 0.0005 | t(38) = 0.785, p = 0.4372 |
| VTA | 16.30007 | 84.49991 | 109.0904 | t(30) = -2.621, p = 0.0204 | t(30) = -2.984, p = 0.0168 | t(30) = -0.791, p = 0.4353 |

**Supplemental Data Table 7 | FOS+ Neuronal Density by ROI and Continuous Behaviors**

| ROI | Huddling Time | | Total Contact | |
|---|---|---|---|---|
| | **Rho** | **P-value** | **Rho** | **P-value** |
| AON | 0.08443115 | 0.7087 | - 0.0599 | 0 .7913 |
| CP | 0.07227838 | 0.7555 | 0.0903 | 0 .6971 |
| NAcc | -0.07256921 | 0.7678 | 0.04212374 | 0.864 |
| NAcsh | 0.05333807 | 0.8233 | 0.1188417 | 0.6178 |
| vBST | 0.1397678 | 0.535 | 0.07003672 | 0.7568 |
| MPOA | 0.472358 | 0.02643 | 0.4817848 | 0.02318 |
| PVT | 0.1834086 | 0.4389 | 0.1940579 | 0.4123 |
| PFC | 0.01026865 | 0.9638 | 0.1875177 | 0.4034 |
| VP | 0.2536314 | 0.2673 | 0.2832088 | 0.2135 |
| PAG | - 0.3196731 | 0.196 | - 0.2013423 | 0.423 |
| VTA | 0.06833592 | 0.7685 | 0.1344592 | 0.5612 |
| BLA | 0.3469362 | 0.1234 | 0.3338747 | 0.1391 |
| LS | - 0.1540298 | 0.4937 | - 0.1084439 | 0.631 |
| LSd | - 0.09527028 | 0.6732 | - 0.04405535 | 0.8457 |

**Supplemental Data Table 8 | FOS+ Neuronal Density in MPOA to Other ROIs by Behavioral Phenotype**

| | All Samples | | Dysparental Samples | | Allopaternal Samples | |
|---|---|---|---|---|---|---|
| ROI | Rho | P-Value | Rho | P-Value | Rho | P-Value |
| AON | 0.3472614 | 0.1133 | 0.5296703 | 0.05142 | 0.2857143 | 0.4927 |
| PFC | 0.5516657 | 0.007777 | 0.7582418 | 0.001673 | 0.1666667 | 0.6932 |
| BLA | 0.2077922 | 0.3661 | -0.1978022 | 0.5171 | 0.7142857 | 0.04653 |
| PAG | 0.08359133 | 0.7416 | 0.5272727 | 0.09557 | -0.2857143 | 0.5345 |
| PVT | 0.4676692 | 0.03759 | 0.7272727 | 0.007355 | -0.02380952 | 0.9554 |
| LS | 0.3980802 | 0.06653 | 0.6879121 | 0.00654 | -0.2142857 | 0.6103 |
| LSd | 0.4929418 | 0.01975 | 0.8153846 | 0.0003791 | -0.2380952 | 0.5702 |
| CP | 0.6103896 | 0.003297 | 0.7967033 | 0.001114 | 0.1666667 | 0.6932 |
| VP | 0.8441558 | 1.507e-06 | 0.8285714 | 0.0002505 | 0.8214286 | 0.02345 |
| NAcc | 0.7263158 | 0.0004292 | 0.8545455 | 0.0008067 | 0.6428571 | 0.08556 |
| NAcsh | 0.7293233 | 0.0002635 | 0.8811189 | 0.0001527 | 0.2857143 | 0.4927 |
| vBST | 0.6668549 | 0.0007002 | 0.832967 | 0.0002166 | 0.1428571 | 0.7358 |
| VTA | 0.4922078 | 0.02341 | 0.6879121 | 0.00654 | 0.07142857 | 0.879 |

**Supplemental Data Table 9 |** Top marker genes for clusters. Clusters ($N$ = 17) are sorted by neurotransmitter type (NT: GABAergic, Glutamatergic, or N/A) and by dominant voltage gated calcium channel (VGCC) subunit. Top marker genes for each cluster (≤ 3) are provided based on differential expression analysis via *FindAllMarkers* in *Seurat*.

| Cluster | NT | VGCC | Top Markers |
|---|---|---|---|
| gab1 | GABA | Cacna2d3 | Slc32a1 |
| gab2 | GABA | Cacna2d3 | Zeb2, Grm7, Foxp2 |
| gab3 | GABA | Cacna2d3 | Celf2, Meis2, Rbfox1 |
| gab4 | GABA | Cacna2d2 | Sox6, Bcl11a, Adarb2 |
| gab5 | GABA | n/a | Ntng1, Pdzrn4 |
| gab6 | GABA | n/a | Rorb, Zpf804b, Zfhx4 |
| gab7 | GABA | Cacna2d2 | Unc5d, Cdh18, Zfhx3 |
| gab8 | GABA | Cacna2d3 | Meis2, Foxp2, Erbb4 |
| gab9 | GABA | Cacna2d2 | Prkca, Megf11, Prdm16 |
| gab10 | GABA | Cacna2d2† | Alk, Pbx3, Adcy2 |
| gab11 | GABA | n/a | Tac2, Grik1, Cadps2 |
| gab12 | GABA | Cacna2d2 | Adarb2, Nrg3, Gria4 |
| glu1 | Gluta | Cacna2d1 | Cacna2d1 |
| glu2 | Gluta | Cacna2d1 | Ebf1, Gpc6, Ralyl |
| glu3 | Gluta | Cacna2d1 | Calcr, Cacna2d1, Il1rapl1 |
| glu4 | Gluta | Cacna2d1 | Cacna2d1, Nxph1, Galntl6 |
| npep | N/A | Cacna2d1 | Rbms3, Pde4b, Caprin2 |

†Also enriched for Cacna1a

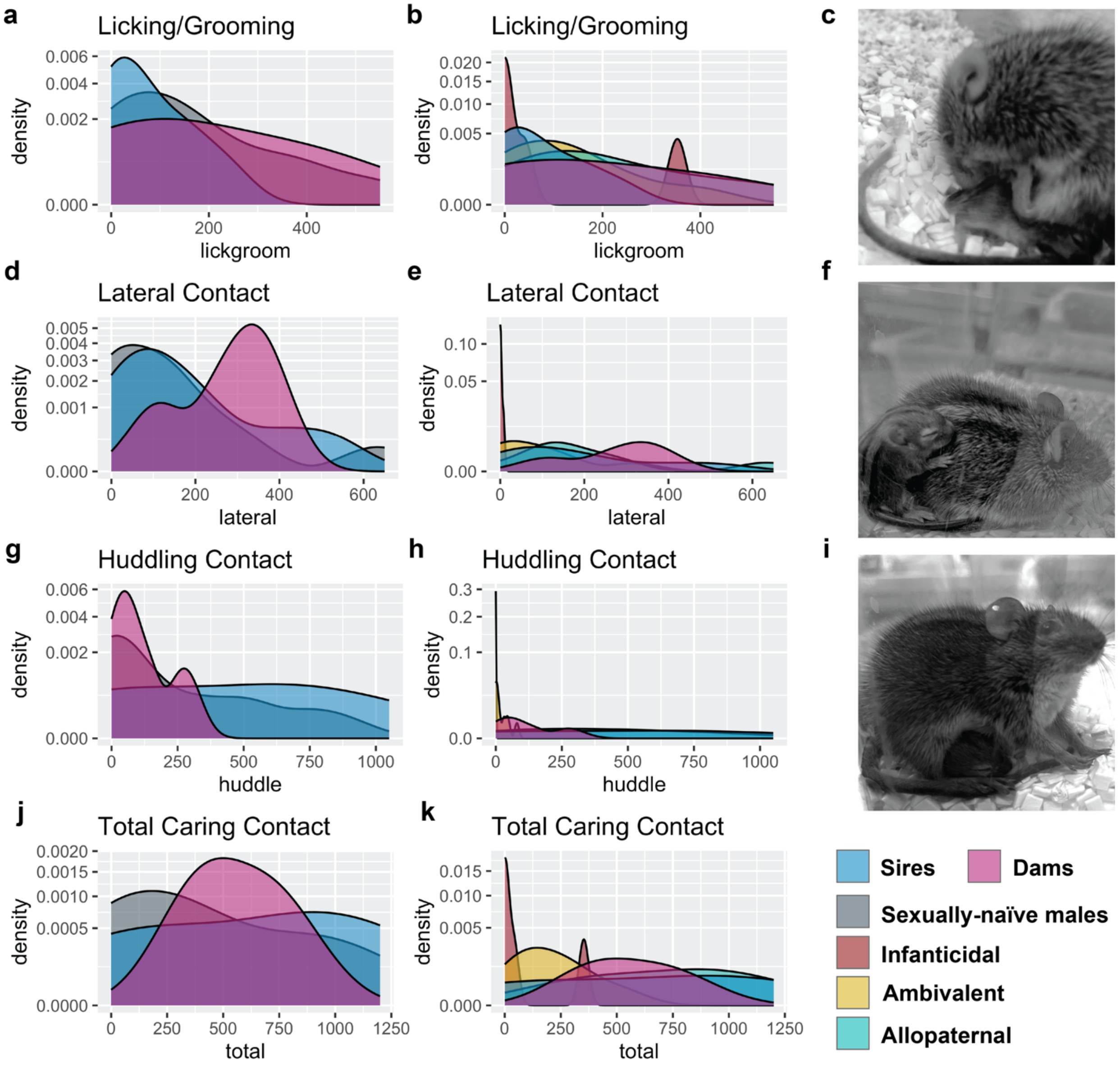


**Supplemental Data Fig. 1 | Distribution of care behaviors in striped mice.**
Density plots illustrating the distribution of individual care behaviors according to sex and reproductive experience alone (**a,d,g,j**) or subdivided by behavioral phenotype (**b,e,h,k**). Sexually-naïve males can be broken down into three groups: Infanticidal, Ambivalent, and Allopaternal. Individual behaviors include licking and grooming (**a-c**), lateral contact (**d-f**), and huddling contact (**g-i**), which cumulatively sum to total caring contact (**j-k**).

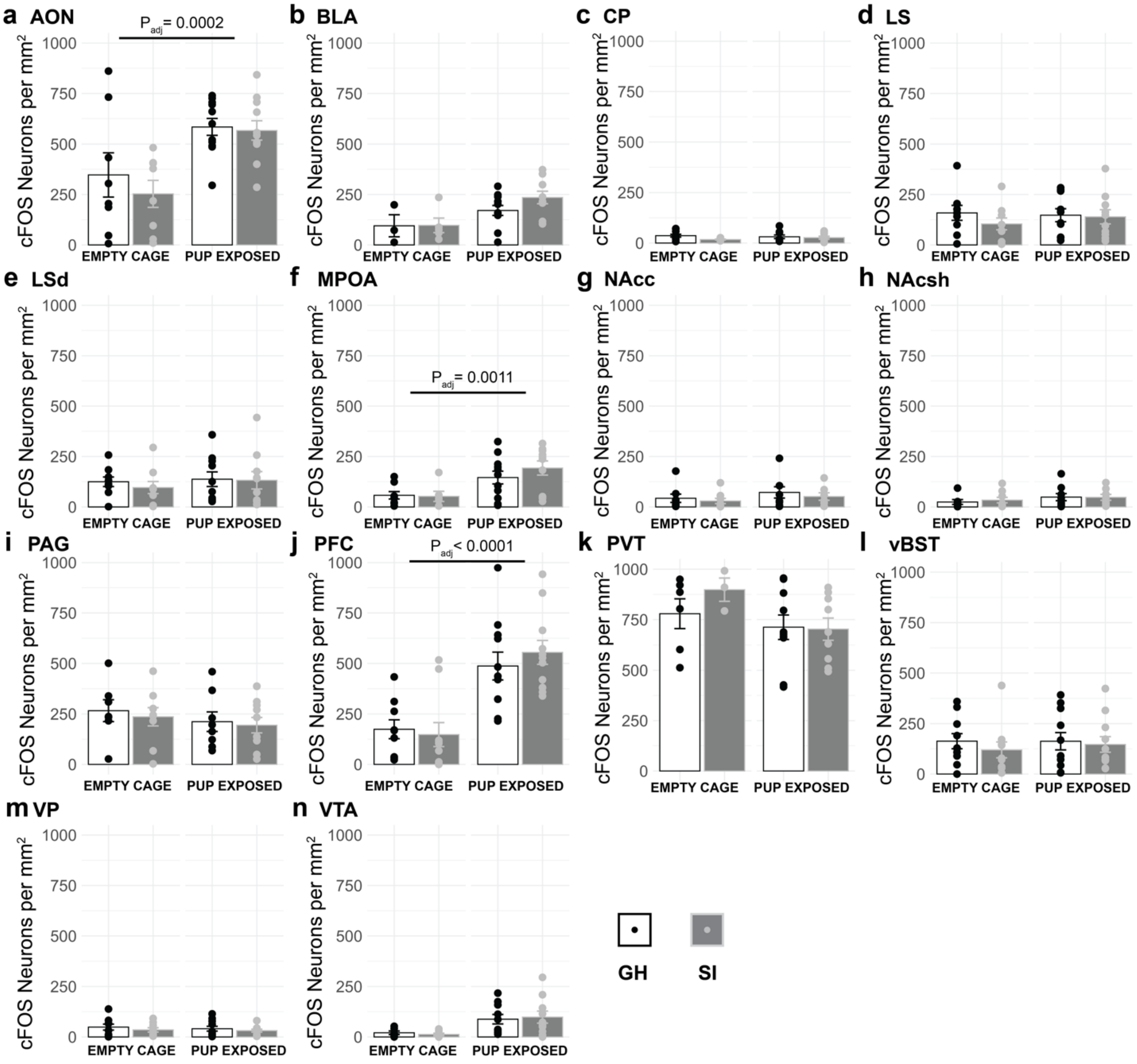


**Supplemental Data Fig. 2| cFOS+ neuron density by pup exposure status & rearing condition**
(**a-n**) The expression of the immediate early gene cFOS was measured via immunohistochemistry in 14 brain regions. Tissue was collected from animals that were either exposed to an empty cage ("empty") or exposed to a pup ("pup) in the hour preceding sacrifice. Animals were either reared in group housing (GH, white bars) or social isolation (SI, grey bars). All p-values are FDR corrected. Error bars represent standard error of means. Abbreviations: (**a**) accessory olfactory nucleus (AON); (**b**) basolateral amygdala (BLA); (**c**) caudate putamen (CP); (**d**), lateral septum (LS); (**e**) dorsal lateral septum (LSd); (**f**) medial preoptic area (MPOA); (**g**) nucleus accumbens core (NAcc); (**h**) nucleus accumens shell (NAcsh); (**i**) periaqueductal grey (PAG); (**j**) prefrontal cortex (PFC); (**k**) periventricular nucleus of the thalamus (PVT); (**l**) ventral bed nucleus of the stria terminalis (vBST); (**m**) ventral pallidum (VP); and (**n**) ventral tegmental area (VTA).

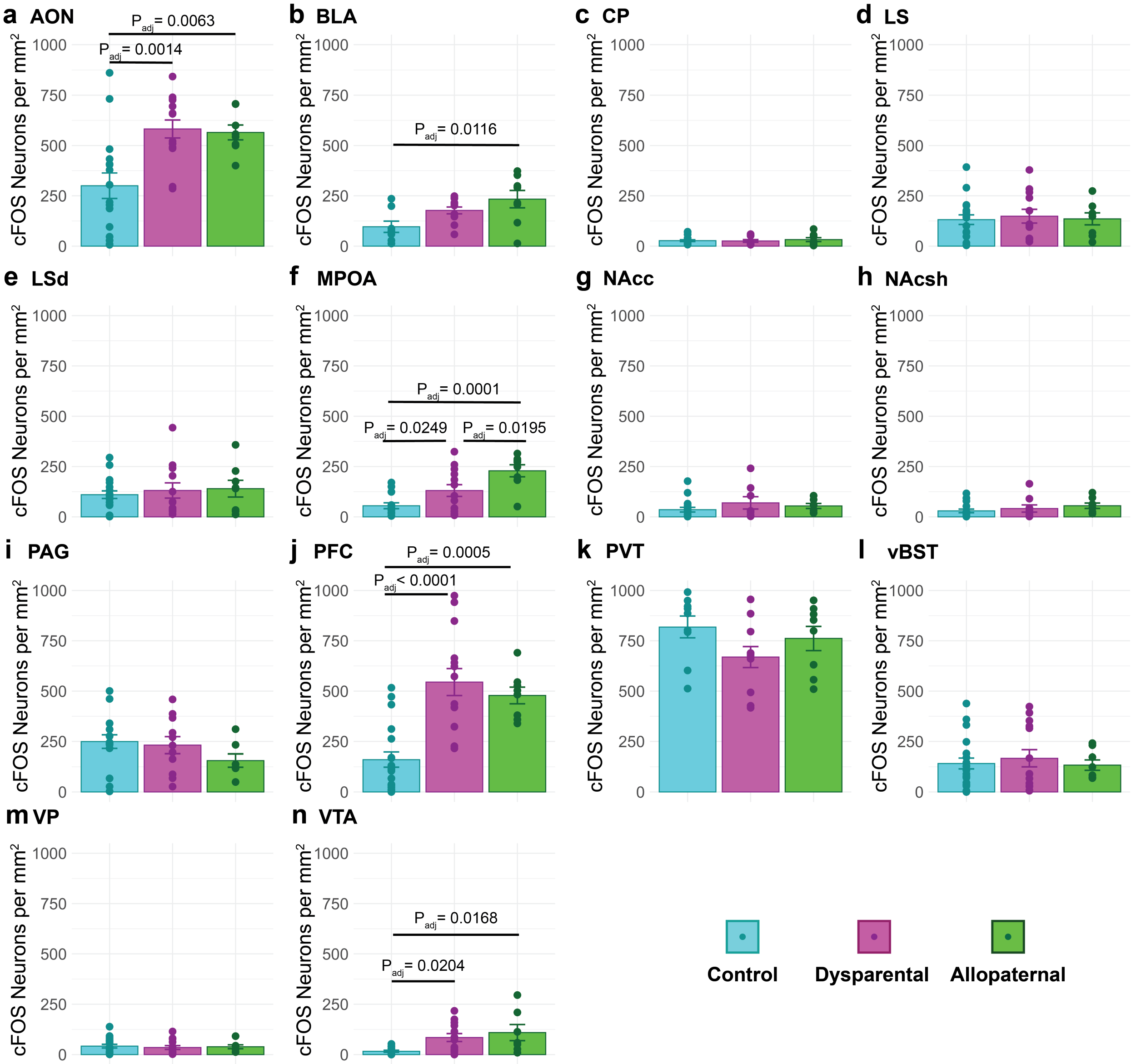


**Supplemental Data Fig. 3 | cFOS+ neuron density by phenotype**

(**a-n**) The expression of the immediate early gene cFOS was measured via immunohistochemistry in 14 brain regions. Tissue was collected from animals who were sorted into three phenotypic groups according to their interactions with pup stimuli: control (empty cage, no pup stimulus, cyan); dysparental (infanticidal and ambivalent, magenta); and allopaternal (green). All p-values are FDR corrected. Error bars represent standard error of means. Abbreviations: (**a**) accessory olfactory nucleus (AON); (**b**) basolateral amygdala (BLA); (**c**) caudate putamen (CP); (**d**), lateral septum (LS); (**e**) dorsal lateral septum (LSd); (**f**) medial preoptic area (MPOA); (**g**) nucleus accumbens core (NAcc); (**h**) nucleus accumens shell (NAcsh); (**i**) periaqueductal grey (PAG); (**j**) prefrontal cortex (PFC); (**k**) periventricular nucleus of the thalamus (PVT); (**l**) ventral bed nucleus of the stria terminalis (vBST); (**m**) ventral pallidum (VP); and (**n**) ventral tegmental area (VTA).

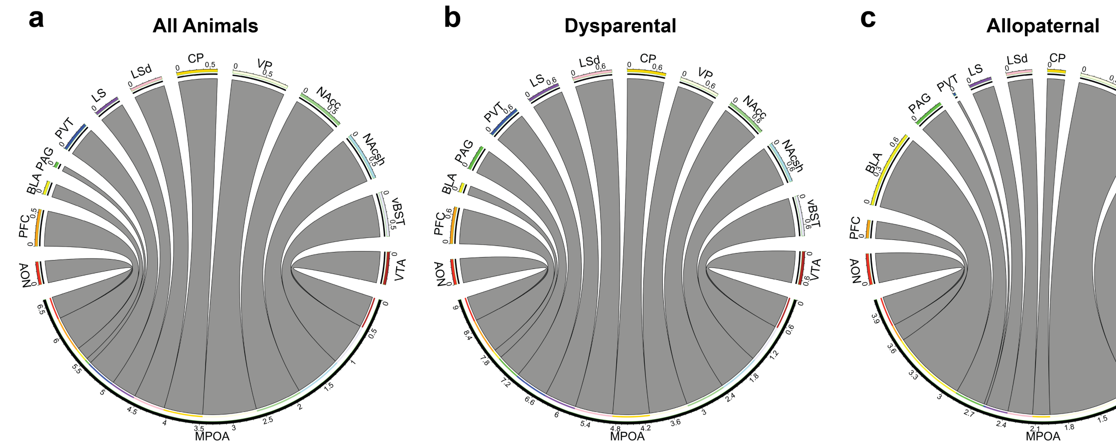


**Supplemental Data Fig. 4 | Correlated activity from the MPOA to other regions of interest**
The pattern of correlations between cFOS expression in the MPOA and 13 other brain regions differs by phenotype: (a) all animals; (b) dysparental animals; (c) allopaternal animals. See Supplemental Data Table 8 for rank correlation coefficients and significance. Abbreviations: accessory olfactory nucleus (AON); basolateral amygdala (BLA); caudate putamen (CP); lateral septum (LS); dorsal lateral septum (LSd); medial preoptic area (MPOA); nucleus accumbens core (NAcc); nucleus accumens shell (NAcsh); periaqueductal grey (PAG); prefrontal cortex (PFC); periventricular nucleus of the thalamus (PVT); ventral bed nucleus of the stria terminalis (vBST); ventral pallidum (VP); and ventral tegmental area (VTA)

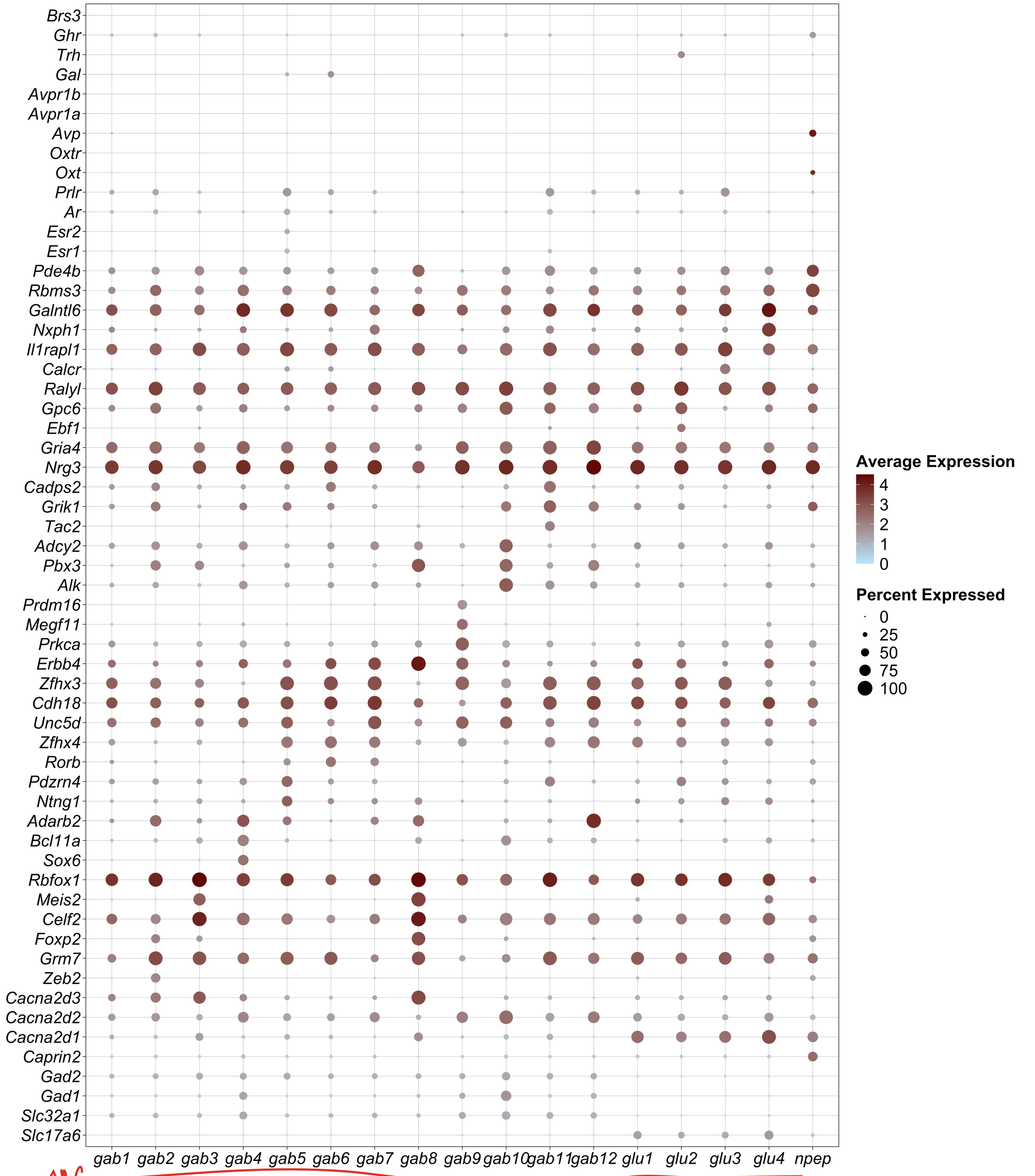


**Supplemental Data Figure 5 |** Dot plot of average expression data by cluster. From bottom to top, marker genes are considered for neurotransmitter type, voltage gated calcium channel subunit, top marker genes for clusters, and neuropeptides of *a priori* interest.

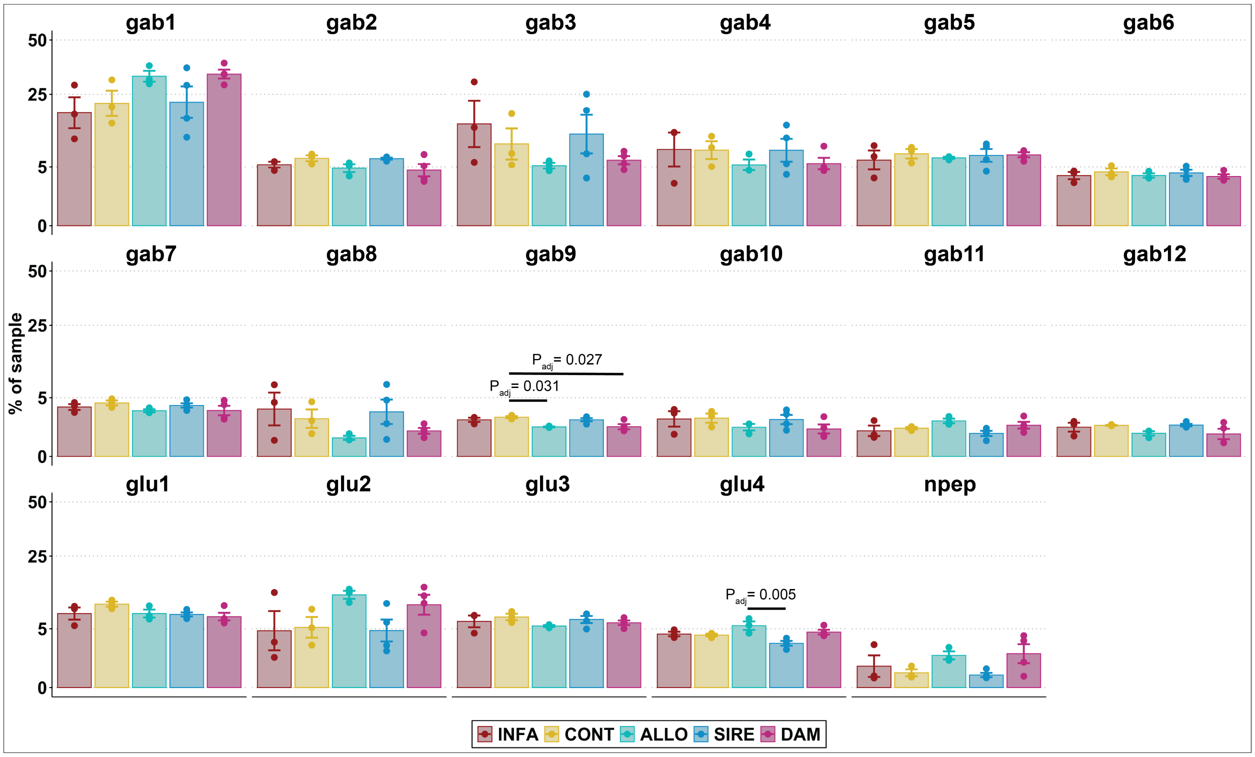


**Supplemental Data Fig. 6 | Percent of sample by neuronal cluster**

Neurons grouped into 17 clusters, including 12 GABAergic (gab1-gab12), four glutamatergic (glu1-4), and one neuropeptidergic cluster (npep). One-way ANOVAs were run to assess whether the relative composition of neurons varied across the five represented groups: infanticidal (INFA), control (CONT), allopaternal (ALLO), sires (SIRE), and dams (DAM). All p-values were generated using Tukey multiple comparisons of means. Error bars represent standard error of means. ANOVA results for cluster 'gab9': ($F(4,12) = 5.3377$, $p = 0.01051$, partial $eta^2 = 0.64$). ANOVA results for cluster 'glu4' ($F(4,12) = 5.475$, $p = 0.009593$, partial $eta^2 = 0.65$).

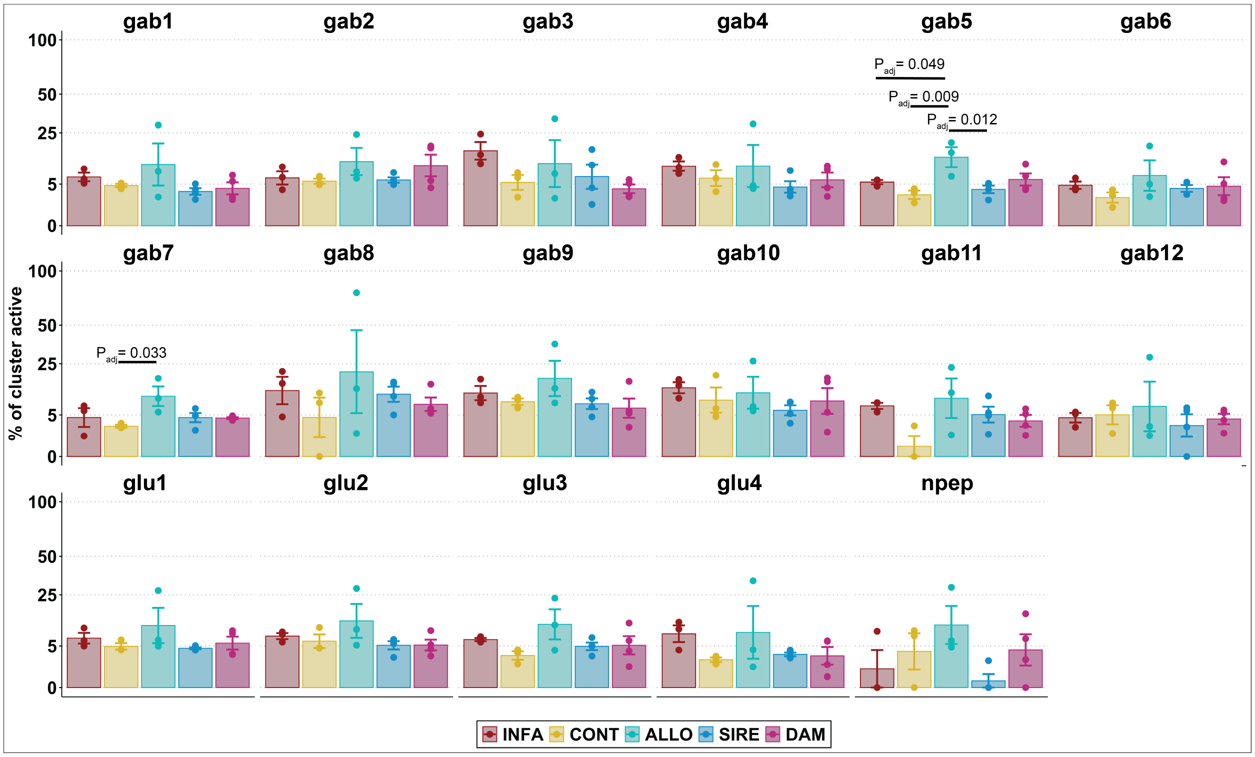


**Supplemental Data Fig. 7 | Percent of respective neuronal clusters with active neurons**

Within each of the 17 neuronal clusters, the number of active neurons was divided by the number of total neurons. One-way ANOVAs were run to assess whether the relative percentage of active neurons varied across the five represented groups: infanticidal (INFA), control (CONT), allopaternal (ALLO), sires (SIRE), and dams (DAM). Active neurons were designated based on expression (> 0 counts) of at least one of 9 immediate early genes (IEGs): *Fos, Fosb, Fosl1, Npas4, Egr1, Egr3, Arc, Jun,* and *Junb.* ANOVA results for 'gab5': (F(4,12) = 5.5665, p = 0.009036, partial eta$^2$ = 0.65). ANOVA results for 'gab7': (F(4,12) = 3.5337, p = 0.03978, partial eta$^2$ = 0.54). All p-values were generated using Tukey multiple comparisons of means. Error bars represent standard error of means.

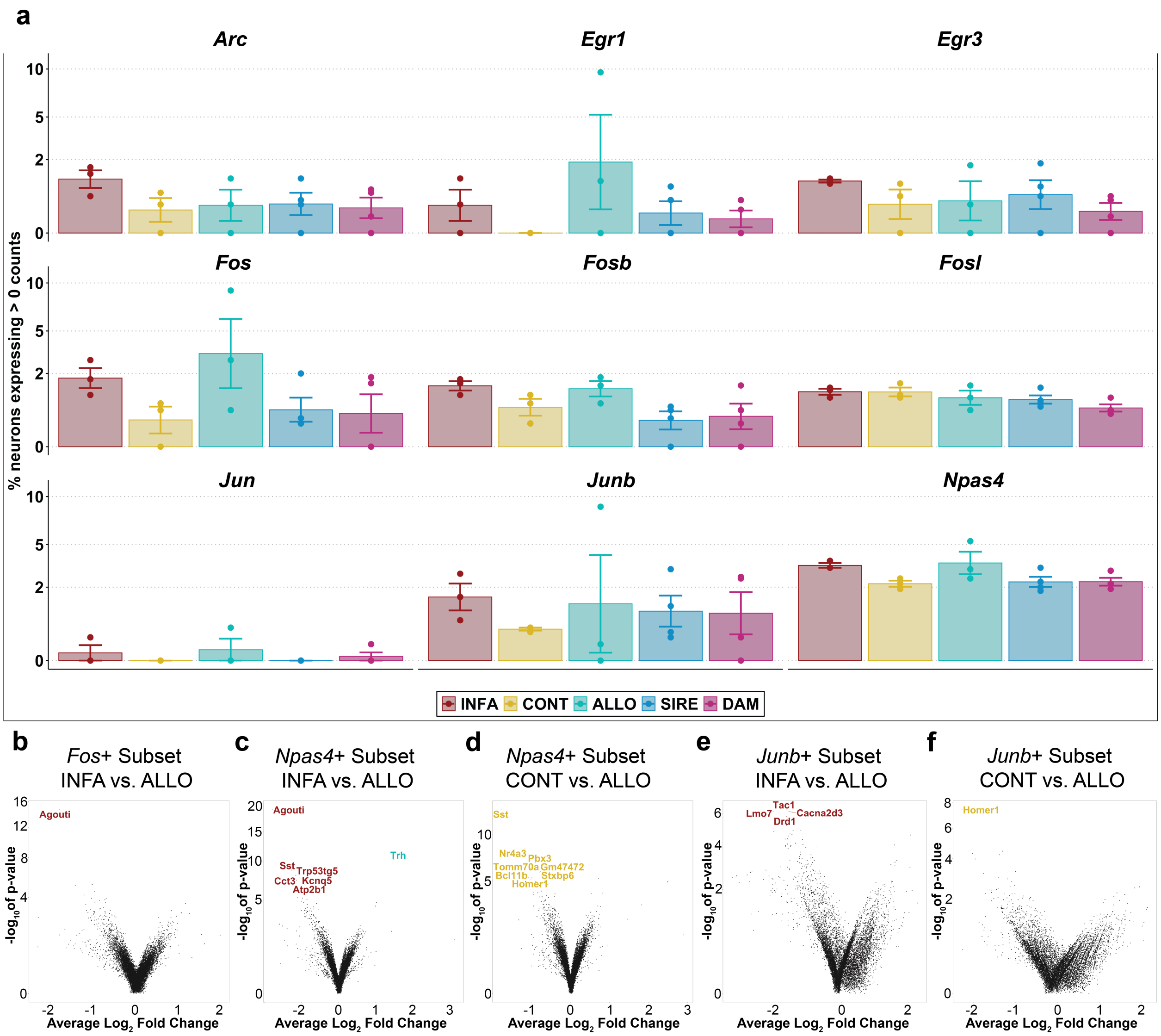


**Supplemental Data Fig. 8 | Immediate early gene expression across neurons**

(**a**) The expression of immediate early genes (IEGs) at the levels of specific IEG and group. Main effects not displayed: *Npas4* was more widely expressed than *Arc* ($p_{adj} = 0.0001921$), Egr1 ($p_{adj} = 0.0022272$), Egr3 ($p_{adj} = 0.0004410$), Fosb ($p_{adj} = 0.0019331$), Fosl1 ($p_{adj} = 0.0044410$), and Jun ($p_{adj} = 0.0000018$). Allopaternal males (ALLO) had wider IEG expression than controls (CONT; $p_{adj} = 0.0060731$), sires (SIRE; $p_{adj} = 0.0109813$), and dams (DAM; $p_{adj} = 0.0064330$), but not infanticidal (INFA; $p_{adj} = 0.4861394$). All p-values were generated using Tukey multiple comparisons of means. Error bars represent standard error of means. (**b-f**) Pseudobulk results within neuronal populations isolated for expression of the most highly expressed IEGs (i.e., *Fos, Npas4,* and *Junb*), comparisons between infanticidal males (left, red), and allopaternal males (right, turquoise); or alloparents and controls (left, gold). Differential gene expression was determined using DESeq2. Named genes were significant after FDR correction (FDR $\leq 0.1$).

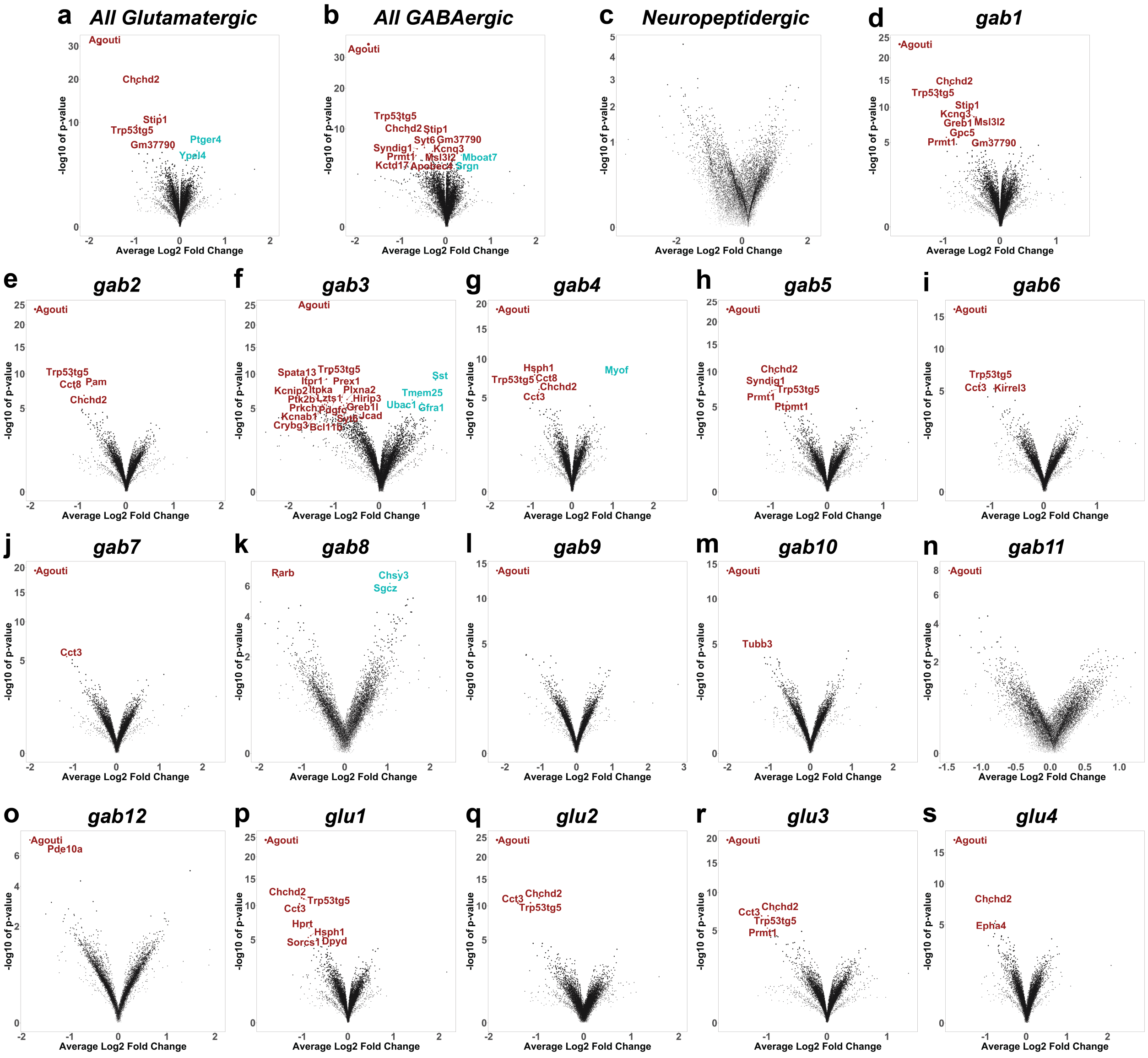


**Supplemental Data Fig. 9 | Pseudobulk DEG analyses in individual neural clusters**
Pseudobulk differential expression analysis (DESeq2) was performed for individual neuronal clusters between infanticidal and allopaternal males and visualized as volcano plots [log2(fold-change) vs -log10(p-adj)]. In all panels, named genes are differentially expressed (FDR ≤ 0.1). Genes in red and to the left are enriched in infanticidal males. Genes in turquoise and to the right are enriched in allopaternal males. (**a**) Glutamatergic neurons were collectively grouped, as were (**b**) GABAergic neurons. (**c-s**) Pseduobulk analyses for smaller clusters, represented in Figure 3, are also presented here.

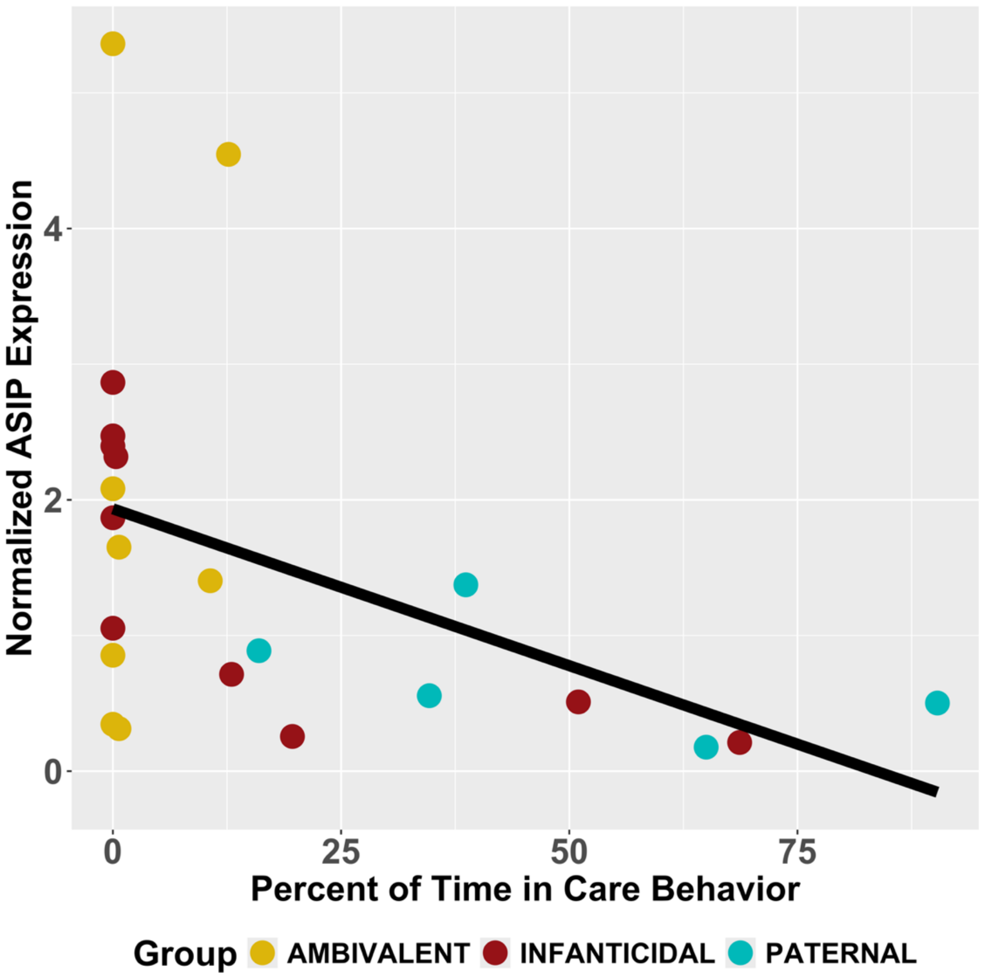


**Supplemental Data Fig. 10 | Validation of correlation between *agouti* expression and behavior**
The correlation between total contact time and *agouti* expression in a sample of sexually naïve group housed males (N = 23, t(21) = -2.3233, p = 0.030, r = -0.45).

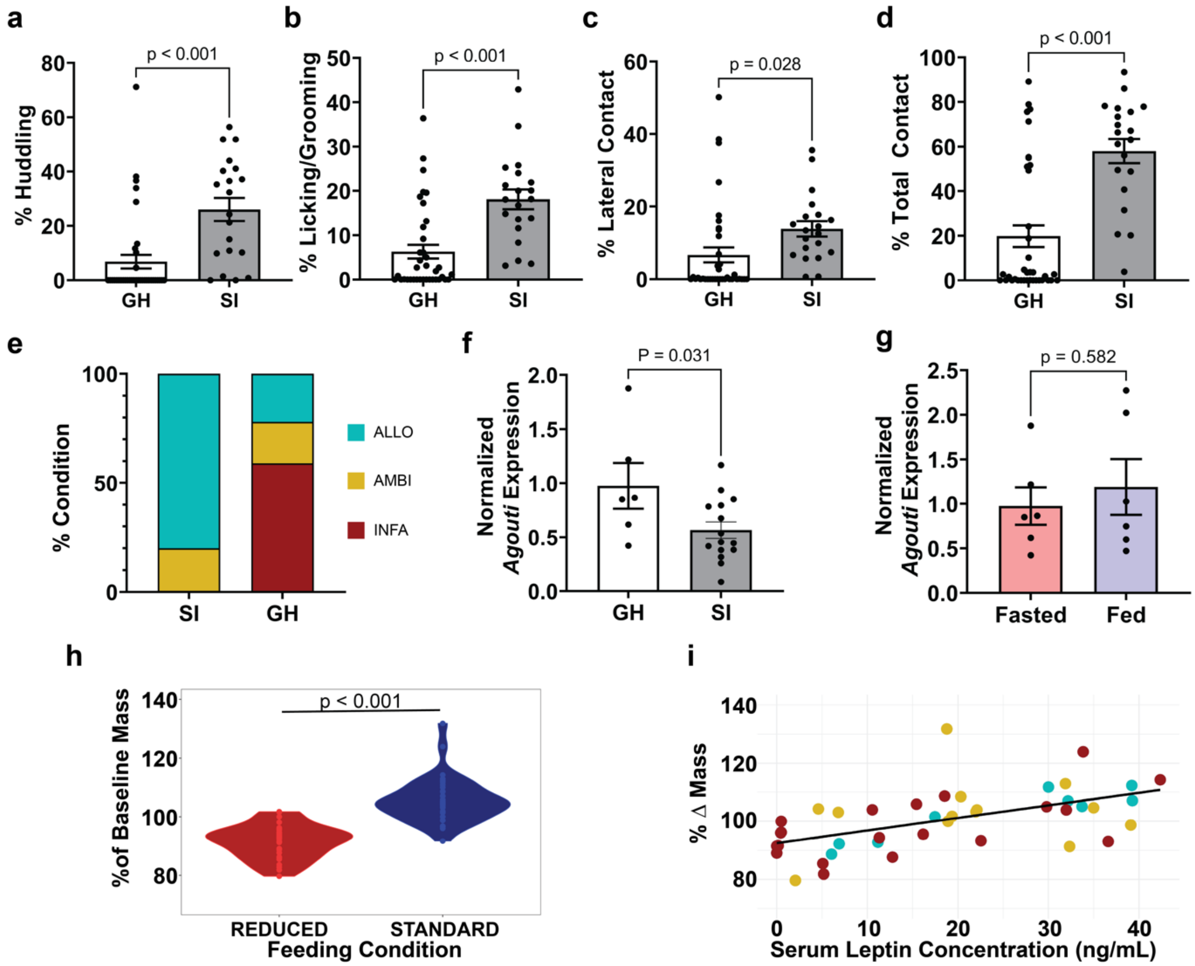


**Supplemental Data Fig. 11 | Baseline allopaternal behavior and *Agouti* expression in fasted males**
Animals were reared in SI and GH conditions. Behavioral outcomes compared at baseline in a fasted state include (**a**) huddling, (**b**) licking/grooming, (**c**) lateral contact, and (**d**) cumulative care. (**e**) The distribution of behavioral phenotypes in SI and GH animals (Chi-square = 22.41, df = 2, $p < 0.0001$). (**f**) MPOA *Agouti* expression in SI and GH males. (**g**) Expression of MPOA *Agouti* compared between animals fasted (i.e., fed ~ 24 hours earlier) or fed (i.e., fed ≤ 2 hours earlier). (**h**) Percent change in weight from pre-test to post-test among males on a reduced diet (n=29; mean of 91.5% of starting weight) vs. a standard diet (n=34; mean of 105% of starting weight), $t(59.867) = -8.282$, $p = 1.647e-11$, $d = -2.04$). (**i**) Serum leptin concentration correlated with change in weight (rho = 0.5961, $p = 3.103e-05$).
